# Identification of a plasma membrane bicarbonate transporter critical for *Leishmania* virulence

**DOI:** 10.1101/2025.07.16.665125

**Authors:** Arunava Seth, Sourav Priyam Adhya, Rupak Datta

**Author notes:** Corresponding author: Rupak Datta.

## Abstract

*Leishmania* parasites survive and proliferate within the acidic phagolysosomes of host macrophages while maintaining a near-neutral cytosolic pH, a process essential for intracellular survival and virulence. Although our prior work implicated a bicarbonate-based buffering mechanism, the absence of any annotated bicarbonate transporter in the parasite genome has hindered its identification. Using an isotope ratio mass spectrometry-based assay, we first confirmed the presence of a functional bicarbonate transport system in the parasite. Guided by this, a noncanonical homology search identified an uncharacterized protein, hereafter designated LmSLC26A, with similarity to metazoan SLC26 family of multi-anion transporters. LmSLC26A localized to the parasite plasma membrane and, under acidic conditions, was found to associate with the membrane carbonic anhydrase LmCA2. Disruption of a single LmSLC26A allele significantly reduced bicarbonate uptake, leading to intracellular acidification that was reversible upon exogenous bicarbonate supplementation. LmSLC26A-deficient parasites exhibited sluggish growth, increased apoptosis, reduced production of virulence associated cAMP and impaired exosome release. Consequently, these mutant parasites showed diminished survival within host macrophages and markedly attenuated virulence in mice. Interestingly, LmSLC26A harbours a unique histidine ecto-phosphatase domain, revealing an unprecedented functional coupling between bicarbonate transport and enzymatic activity. Together, these findings establish LmSLC26A as the first identified bicarbonate transporter not only in *Leishmania* but also across the broader trypanosomatid and protozoan lineages, and define bicarbonate transport as a critical determinant of pH homeostasis and pathogenicity in the parasite.

## Introduction

*Leishmania* species are kinetoplastid protozoan parasites of the trypanosomatid family that cause leishmaniasis, a group of neglected tropical diseases affecting millions of people worldwide. The clinical manifestations of leishmaniasis vary widely depending on the infecting species, ranging from localized cutaneous lesions to life-threatening visceral disease (1). Current treatment options remain limited due to drug toxicity, high cost, and the increasing emergence of resistant strains, highlighting the need for a deeper understanding of parasite biology to identify new therapeutic targets (2). During its digenetic life cycle, *Leishmania* parasites alternate between a slender, flagellated promastigote stage in the sandfly vector and a rounded, non-flagellated amastigotes within the phagolysosomes of mammalian macrophages (3). Successful adaptation to these distinct environments requires remarkable physiological flexibility, particularly during the transition from the near-neutral pH of the sandfly midgut to the highly acidic milieu of the host phagolysosome. Notably, both promastigote and amastigote forms are able to maintain a near-neutral intracellular pH even when exposed to external conditions as acidic as pH ∼5.0, closely resembling the phagolysosomal environment (4). This capability is especially striking because the parasite exploits the resulting transmembrane proton gradient to drive H^+^-coupled uptake of essential nutrients such as glucose and amino acids, a process that would typically lead to cytosolic acidification. How *Leishmania* maintains cytosolic pH homeostasis while thriving in such acidic conditions remains a longstanding and incompletely understood question.

Cytosolic pH homeostasis in *Leishmania* was previously believed to depend exclusively through active proton extrusion mediated by membrane-associated P-type H^+^-ATPases (5). However, our recent findings revealed a more energy-efficient mechanism in *Leishmania major* involving the coordinated action of two carbonic anhydrases (CAs), LmCA1 in the cytosol and LmCA2 at the plasma membrane that together maintain pH balance and promote acid tolerance in this parasite. Interestingly, when exposed to an acidic medium, haploinsufficient mutants of LmCAs (LmCA1^+/-^, LmCA2^+/-^, LmCA1^+/-^ :LmCA2^+/-^) exhibited intracellular acidosis and growth retardation, both of which could be prevented by exogenous bicarbonate supplementation. Based on these findings, it was proposed that LmCA1 buffers intracellular protons by converting them into water and diffusible CO_2_, which exits the cell. At the plasma membrane, LmCA2 hydrates this CO_2_ to produce extracellular bicarbonate and protons. To sustain this cycle and maintain cytosolic buffering capacity, extracellular bicarbonate must be transported back to the cell, since it is not freely permeable across membranes. This led to the hypothesis of a dedicated bicarbonate transporter, likely functioning in concert with membrane-localized LmCA2, to efficiently preserve intracellular pH neutrality, particularly under acidic stress (6). A key role of bicarbonate transport in maintaining pH balance in *L. major* was also supported by the findings of Vieira *et al.*, who showed that exogenous addition of bicarbonate reverses acidification caused by H^+^ pump inhibition and elevates the steady state intracellular pH even in untreated promastigotes (7). In the present study, we also provide direct experimental evidence for bicarbonate transport in *L. major* cells using a newly established isotope ratio mass spectrometry (IRMS) based assay. However, these biochemical evidences, which strongly indicated the presence of a functional bicarbonate transporter in *Leishmania*, appeared contradictory to the absence of any annotated bicarbonate transporter in the parasite genome (8). Thus, uncovering the molecular identity of the bicarbonate transporter in *Leishmania* posed a significant challenge.

While bicarbonate transporters remain unexplored in protozoan parasites, with none functionally validated in *Leishmania* or any other member of the Kinetoplastida phylum, they are fairly well-characterized in metazoans, where they primarily belong to two major families: SLC4 and SLC26 (9, 10). The SLC4 family of transporters, which typically contain 10 - 14 transmembrane domains, mediate bicarbonate transport across the plasma membrane with either chloride or sodium ions. They play critical roles in intracellular pH regulation, CO_2_ transport in erythrocytes, H^+^/HCO ^-^ exchange in epithelial tissues, and in controlling cell volume (9). In contrast, members of the SLC26 family act as versatile multi-anion exchangers, transporting bicarbonate, chloride, sulfate, oxalate or other anions. Like SLC4 proteins, they contain 10 - 14 transmembrane domains, but are uniquely defined by a conserved C-terminal STAS (sulfate transporter and anti-sigma factor antagonist) domain, which regulates their stability, intracellular trafficking, and interactions with partner proteins. These transporters contribute to a range of physiological processes, including transepithelial Na^+^ and Cl^-^ movement, bicarbonate reabsorption in the nephron, bicarbonate secretion by the exocrine pancreas and regulation of pulmonary ion balance (11, 12). Importantly, mutations in several members of SLC26 family underlie human diseases such as hereditary deafness, chondrodysplasias, chloride-losing diarrhea and Pendred syndrome (13).

Since our initial analysis of the *Leishmania* genome failed to identify any classical bicarbonate transporter, we adopted an alternative strategy and searched for non-canonical candidates with potential bicarbonate transport activity. Through this approach, we identified the sequence of a 1982 - amino acid uncharacterized protein (gene ID: LmjF.28.1690), annotated as ‘sulfate transporter-like protein’ in the *L. major* genome. Based on its strong homology with human SLC26A3 and the presence of the characteristic STAS domain, we designated this protein as LmSLC26A. While LmSLC26A retains conservation of key functional residues with human SLC26A3, particularly within the transmembrane and STAS domains, it also displays novel features including a histidine phosphatase domain, an extended C terminal tail, and an unusually large overall size compared with canonical SLC26 members, suggesting evolutionary divergence. Interestingly, in some prokaryotes, such as marine cyanobacteria, SLC26-related sulfate permeases (SulP) function as bicarbonate transporters, serving as major pathways for inorganic carbon entry and contributing substantially to global CO_2_ sequestration (14). Could LmSLC26A represent the elusive bicarbonate transporter in *Leishmania*, we asked.

By combining biochemical analyses with genetic perturbation approaches, we provide strong evidence that LmSLC26A functions as a bicarbonate transporter, the first identified not only in *Leishmania* but also across the broader trypanosomatid and protozoan lineages. We demonstrate that LmSLC26A localizes to the parasite plasma membrane, where it plays a critical role in maintaining cytosolic pH homeostasis, supporting parasite growth, and promoting survival under acidic conditions. Loss of LmSLC26A triggered apoptosis, disrupted cAMP production, and markedly reduced the release of parasite-derived exosomal vesicles. Consistent with these cellular defects, LmSLC26A-deficient parasites exhibited impaired intracellular survival in macrophages and severely attenuated virulence in mice. Together, these findings identify LmSLC26A-mediated bicarbonate transport as a key determinant of acid acclimation and pathogenicity in *Leishmania* and establish it as a potential therapeutic target.

## Results

### *Leishmania* possesses a functional bicarbonate transport system

To determine whether *L. major* cells are sensitive to inhibition of bicarbonate transport, we treated them with DIDS (4,4′-diisothiocyanatostilbene-2,2′-disulfonate), a well-established inhibitor of chloride-bicarbonate exchangers (9). DIDS caused a dose-dependent reduction in cell growth under normal conditions (pH 7.2), which was further diminished at acidic pH (5.5), mimicking the phagolysosomal environment. Supplementation with extracellular bicarbonate partially restored growth under both conditions, suggesting that the involvement of bicarbonate transport in observed growth defect (Fig. 1 A and B). To directly assess bicarbonate uptake, we developed an isotope ratio mass spectrometry (IRMS)-based assay. *L. major* cells were incubated with isotopically labeled ^13^C-bicarbonate, and its intracellular accumulation was quantified by measuring the δ^13^C values using isotope ratio mass spectrometry. The δ^13^C values were reported in per mille (‰) relative to the Vienna Pee Dee Belemnite (VPDB) standard, a widely used reference in stable isotope analysis that enables accurate comparison of isotopic values across samples (15). Most biological samples exhibit negative δ^13^C values because they contain relatively lower proportions of ^13^C atoms compared to ^12^C than the VPDB standard, typically ranging between -35‰ and -5‰ (16). As per the experimental plan, intracellular accumulation of ^13^C-bicarbonate would be reflected by an increase in cellular δ^13^C values, indicating bicarbonate uptake into the cells (Fig. 1C). To examine the time dependence of bicarbonate uptake, *L. major* promastigotes were incubated with ^13^C-bicarbonate for 2.5, 5 and 10 min. Cellular δ13C values increased progressively by 2.2‰, 7.1‰ and 9.7‰ at 2.5, 5 and 10 min, respectively, corresponding to ∼1.1, 1.6 and 2.1-fold enrichment over the basal level. These findings demonstrate time-dependent accumulation of bicarbonate-derived ^13^C in *Leishmania* cells (Fig. 1D). Because bicarbonate is a charged molecule that cannot freely diffuse across the lipid bilayer, these results strongly suggest the presence of a functional transmembrane bicarbonate transport system that is essential for parasite physiology.

**Figure 1.**
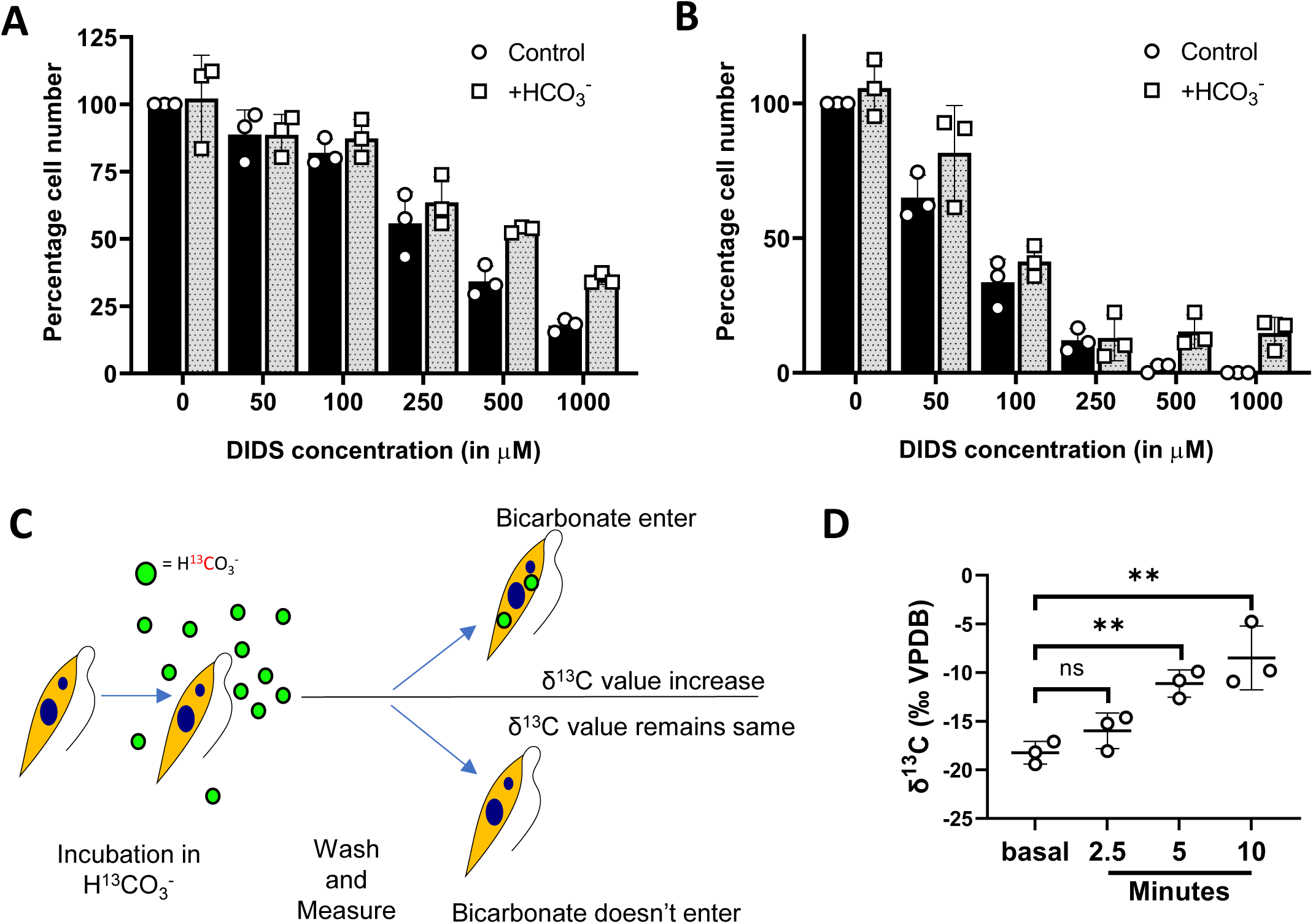
Evidence supporting the presence of a functional bicarbonate transport system in *Leishmania*. (A) *L. major* promastigotes were grown at pH 7.2 or (B) pH 5.5 in the presence of increasing concentrations of the bicarbonate transport inhibitor DIDS, supplemented with 5.9 mM bicarbonate (dotted bar) or without bicarbonate supplementation (black bar). Cells were counted after 72 hours of growth. Error bars represent SD from three independent experiments. (C) Schematic diagram illustrating the principle of the bicarbonate uptake assay based on cellular δ^13^C (relative abundance of the stable carbon isotope ^13^C compared to ^12^C in sample to that of the standard) enrichment following incubation with ^13^C-bicarbonate. (D) δ^13^C values (expressed in ‰ unit against VPDB scale) in *L. major* cells under basal conditions and after 2.5, 5 and 10 minutes incubation in H^13^CO ^-^-containing PBS, measured by IRMS. Error bars represent SD from three independent experiments. ns, P= 0.1467 (between basal and cells at 2.5 minutes of incubation). **, P= 0.0025 (between basal and cells at 5 minutes of incubation). **P = 0.0084 (between basal and cells at 10 minutes of incubation), determined by Student’s t-test.

### Identification of LmSLC26A as the putative bicarbonate transporter in *Leishmania*

Despite strong biochemical evidence for a functional bicarbonate transporter in *Leishmania*, no annotated homolog is present in its genome. We therefore searched for noncanonical candidates, which led us to an uncharacterized 1982-amino acid protein (LmjF.28.1690), annotated in UniProt as a “sulfate transporter-like protein”. NCBI BLAST analysis revealed that LmjF.28.1690 shares strong sequence similarity with human SLC26 family members but not with those of the SLC4 family, prompting us to designate it as LmSLC26A (Fig. 2A, Table S1). Members of the SLC26 family are multifunctional anion transporters capable of mediating bicarbonate exchange (17). Interestingly, comparative analysis showed that LmSLC26A is nearly twice the size of typical SLC26 transporters (Fig. 2A). To check the expression of LmSLC26A, total RNA isolated from *L. major* promastigotes was subjected to RT-PCR. A ∼5.9 kb amplicon corresponding to the full-length LmSLC26A mRNA was obtained, confirming that the gene is actively transcribed (Fig. 2B). Domain analysis using InterPro revealed that LmSLC26A shares key features with human SLC26A3, including eleven transmembrane domains and the STAS domain, which is known to be involved in transporter regulation, trafficking, and protein-protein interactions (11, 18, 19). However, LmSLC26A also exhibits several distinctive features suggestive of evolutionary divergence. In addition to an unusually large overall size and an extended C terminal tail, InterPro analysis predicted the presence of a histidine phosphatase domain spanning residue 730 to 999 (Fig. 2C). Sequence comparison with representative human, bacterial, and fungal histidine phosphatases showed that this region retains the conserved RHGXRXP catalytic motif, characteristic of the histidine phosphatase superfamily (20). The primary catalytic histidine residue is preserved, and most residues within the motif are also conserved in LmSLC26A. However, the second arginine in the canonical RHGXRXP motif is replaced by valine in LmSLC26A (Fig. S1). The presence of an extracellular histidine phosphatase domain in LmSLC26A is unprecedented among characterized bicarbonate transporters, highlighting the uniqueness of this protein. This unusual feature suggests that LmSLC26A may possess additional regulatory or enzymatic capabilities beyond its predicted transporter function. Alignment of the N-terminal transmembrane region of LmSLC26A (residues 1-724) with human SLC26A3 revealed 24% identity and 46% overall similarity, with several residues associated with congenital chloride diarrhea conserved in the *Leishmania* ortholog (Fig. 2D) (18, 21). Based on these bioinformatics analyses, we hypothesized that LmSLC26A could function as a bicarbonate transporter in *Leishmania*.

**Figure 2.**
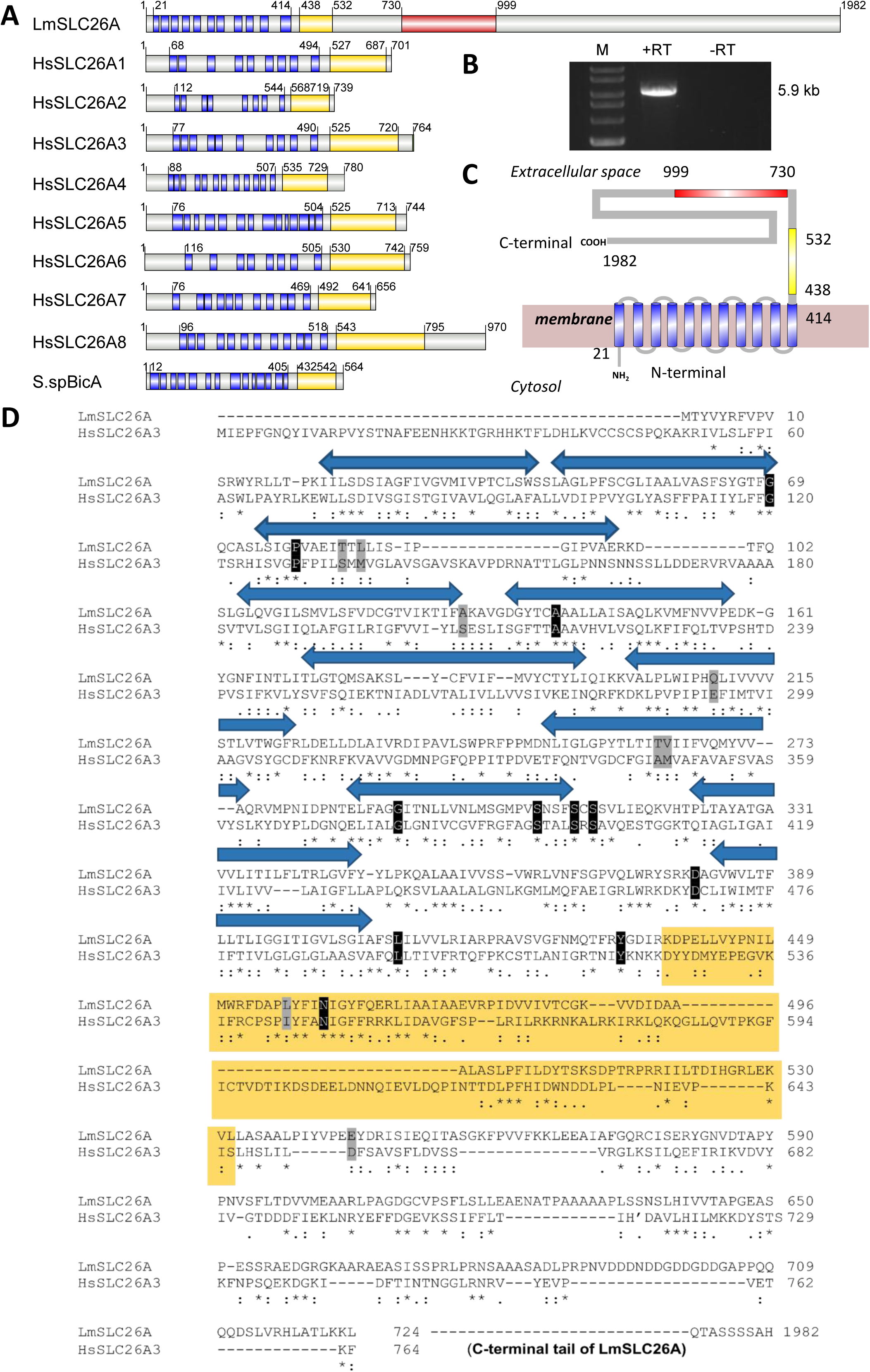
Identification of LmSLC26A as the putative bicarbonate transporter in *Leishmania*. (A) Schematic representation of the domain organization of LmSLC26A in comparison with human SLC26 family members (HsSLC26A1-A8) and the bacterial homolog from Synechococcus sp. (S.spBicA). Predicted transmembrane helices are depicted as blue boxes. The STAS (sulfate transporter and anti-sigma factor antagonist) domains are shown in yellow, while the predicted histidine phosphatase-like domain in LmSLC26A is marked in red. Numbers indicate amino acid positions. (B) Expression analysis of LmSLC26A was performed by semi-quantitative RT-PCR using gene-specific primers (P1/P2). The 5.9 kb amplified product is indicated in the figure (lane marked ‘+’). The lane marked ‘−’ represent negative control reactions lacking reverse transcriptase. (C) Schematic diagram showing the predicted membrane topology of LmSLC26A. The protein is organized into 11 transmembrane (TM) helices (blue cylinders) embedded in the lipid bilayer (light pink), with a cytoplasmic N-terminus and an extended extracellular C-terminal tail (residues 414-1982). The STAS domain (residues 438-582) is indicated in yellow, while the predicted histidine phosphatase domain (residues 740-999) is highlighted in red. (D) Sequence alignment of LmSLC26A^1-764^ with human SLC26A3 (HsSLC26A3). The blue double-sided arrow shows the predicted transmembrane domains. The putative STAS domain (residues 438-582) is marked by yellow boxes. The disease (CCD) associated amino acids in the SLC26A3 gene that are either conserved or are replaced with similar amino acids in LmSLC26A are marked by black or grey boxes, respectively.

### LmSLC26A is localized at the parasite plasma membrane

To determine the subcellular localization of LmSLC26A, we generated antibodies against a C-terminal fragment of the protein encompassing amino acids 597-935 (Fig. S2A). The fragment was cloned into the pET-28a vector, expressed in *E. coli* BL21(DE3), and the recombinant protein was purified (Fig. S2B and C). The purified protein fragment was used to raise polyclonal antibodies in mouse and rabbit and subsequently validated by western blotting against purified antigen (Fig. S2D and E). To further validate the specificity of the antibody for immunofluorescence experiments, we established stable HEK-293 cells expressing LmSLC26A (1–1024), which encompasses the predicted transmembrane region, STAS domain, and histidine phosphatase domain, including the region used for antibody generation (Fig. S3A). The corresponding fragment was cloned into the pCXN vector, and the construct was confirmed by EcoRI restriction digestion (Fig. S3B). The verified construct was subsequently used to generate stable HEK-293 transfectants through G418 selection. Immunofluorescence analysis using the anti-LmSLC26A antibody revealed a detectable signal in LmSLC26A^1-1024^-expressing cells, whereas no signal was detected in control HEK-293 cells (Fig. S3C), confirming the specificity of the antibody for LmSLC26A. Using this anti-LmSLC26A antibody, we performed immunofluorescence in non-permeabilized *L. major* promastigotes (Fig. 3A). Under these conditions, antibodies can access extracellular epitopes but cannot reach intracellular targets when the plasma membrane remains intact. The LmSLC26A-specific signal was clearly detected at the cell periphery, consistent with plasma membrane localization (Fig. 3B). Importantly, the intracellular mitochondrial malic enzyme of *L. major* (LmME) was detected only after cell permeabilization and not under non-permeabilized conditions (Fig. S4). This control demonstrates that intracellular antigens were inaccessible to antibodies under the non-permeabilized conditions, supporting the interpretation that the peripheral LmSLC26A staining represents a surface-accessible pool of the protein. We next examined the localization of LmSLC26A in the intracellular amastigote stage of the parasite. Accordingly, BALB/c mice were infected with *L. major*, and amastigotes were isolated from infected footpad lesions at 12 weeks post-infection (p.i.) for immunofluorescence analysis. A clear membrane-associated signal for LmSLC26A was again observed, indicating that the protein is expressed and correctly targeted to the plasma membrane in the *Leishmania* amastigotes as well (Fig. 3C). Although LmSLC26A is localized at the plasma membrane in both stages of the parasite, a closer examination revealed a difference in its distribution pattern. In promastigotes, LmSLC26A appears in discrete patches along the plasma membrane, whereas in amastigotes it shows more uniform distribution (Fig. 3B and 3C, insets). The functional significance of this difference in membrane distribution, however, remains unclear.

**Figure 3.**
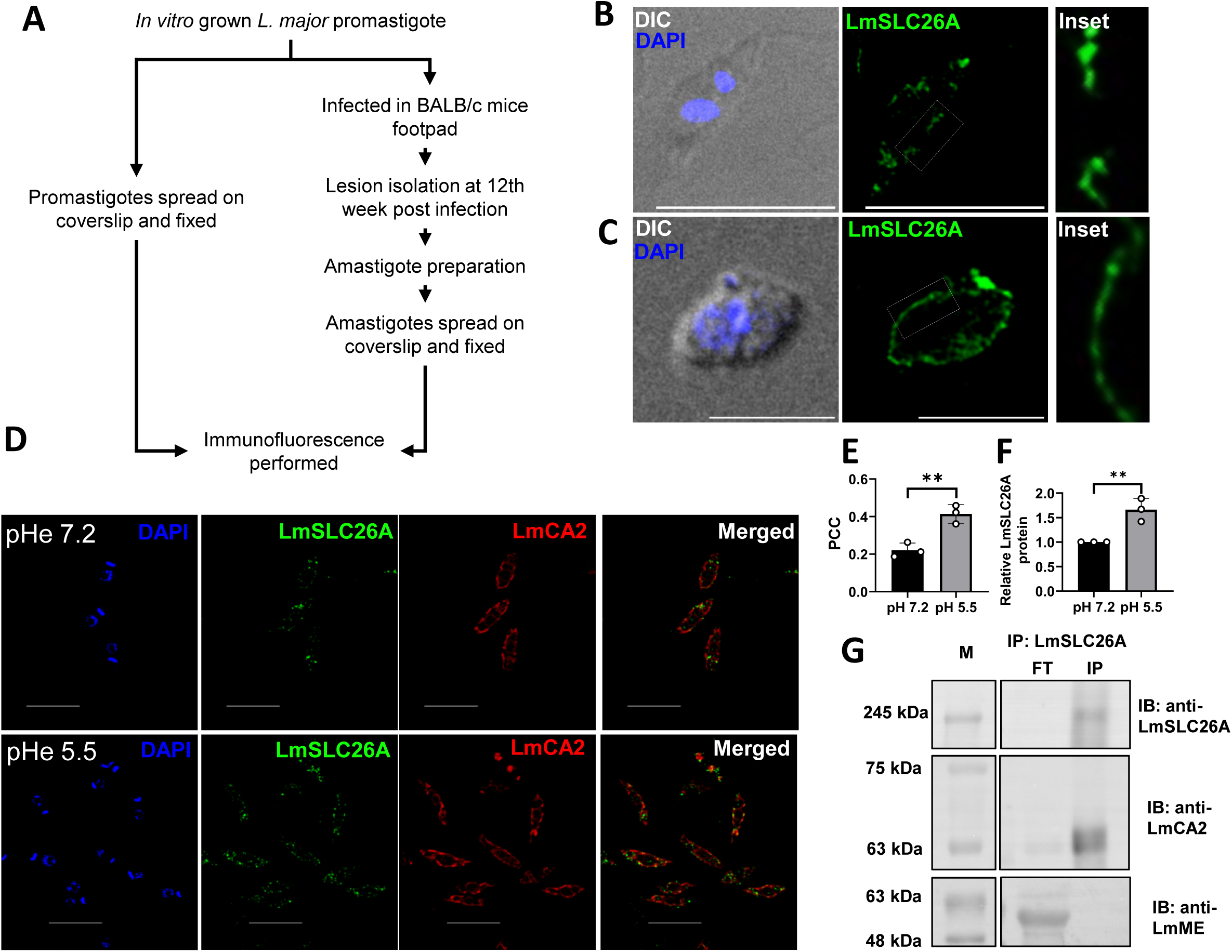
LmSLC26A localizes to the parasite plasma membrane and associates with LmCA2 under acidic conditions. (A) Schematic representation of the workflow used to obtain *L. major* promastigotes and amastigotes for immunofluorescence analysis. (B) *L. major* promastigotes were immunostained with anti-LmSLC26A (green) without permeabilization. DAPI (blue) marks the nucleus and kinetoplast. Cells were imaged using a Leica SP8 confocal microscope. Insets show magnified views of boxed regions. Promastigotes display a discontinuous distribution of LmSLC26A on the plasma membrane. Scale bar: 10 μm. (C) *L. major* amastigotes isolated from infected mouse footpads were immunostained with anti-LmSLC26A (green) without permeabilization. DAPI (blue) marks the nucleus and kinetoplast. Cells were imaged using a Leica SP8 confocal microscope. Insets show magnified views of boxed regions. Amastigotes exhibit a continuous distribution of LmSLC26A on the plasma membrane. Scale bar: 5 μm. (D) *L. major* promastigotes grown at pH 7.2 (top row) or pH 5.5 (bottom row) were co-immunostained with anti-LmSLC26A (green) and anti-LmCA2 (red). DAPI (blue) marks the nucleus and kinetoplast. Merged images indicate colocalization. Cells were imaged using a Leica SP8 confocal microscope. Scale bar: 10 μm. (E) Pearson’s colocalization coefficient was calculated using the JaCoP plugin (ImageJ). At least 60 cells were analyzed per condition. Error bars represent SD from three independent experiments. **P = 0.006 (Student’s t-test). (F) Relative LmSLC26A protein levels under acidic conditions (pH 5.5) compared to neutral conditions (pH 7.2), quantified as mean fluorescence intensity using ImageJ. At least 60 cells were analyzed per condition. Error bars represent SD from three independent experiments. **P = 0.0083 (Student’s t-test). (G) Co-immunoprecipitation of LmCA2 using LmSLC26A as bait. Western blot analysis revealed bands at approximately 215 kDa and 63 kDa, corresponding to LmSLC26A and LmCA2, respectively. An additional western blot using anti-LmME antibody, which recognizes an unrelated parasite protein, showed that LmME was not detected in the LmSLC26A immunoprecipitated fraction. M, molecular weight marker; FT, flow-through; IP, immunoprecipitated fraction; IB, antibody used for immunoblotting.

In human cells, bicarbonate transporters often associate with membrane bound carbonic anhydrases, e.g. CAIV or CAIX, to form a bicarbonate transport metabolon (22, 23). This prompted us to examine whether LmSLC26A interacts with the membrane associated carbonic anhydrase of *L. major*, LmCA2. For this, antibodies were generated against a fragment (amino acids 291-540) of LmCA2 (Fig. S2 F-H). Co-immunofluorescence was then performed with LmSLC26A and LmCA2 antibodies in non-permeabilized *L. major* promastigotes grown under normal (pH 7.2) or acidic (pH 5.5) conditions. Despite both proteins exhibiting typical membrane localization, under normal conditions only a weak co-localization was observed, with a Pearson correlation coefficient (PCC) of 0.23. In contrast, exposure to acidic pH increased the spatial association between LmSLC26A and LmCA2, with the PCC rising to 0.41. This moderate increase in PCC coincided with elevated LmSLC26A signal intensity (Fig. 3D-F), suggesting that acidic conditions may influence the spatial distribution of LmSLC26A relative to LmCA2. To assess whether the observed colocalization reflects a direct physical interaction, co-immunoprecipitation (Co-IP) was performed. Promastigotes grown under acidic conditions were detergent-solubilized, and the proteins in the lysate were immunoprecipitated using anti-LmSLC26A antibody. Interestingly, western blot analysis of the immunoprecipitate confirmed the presence of LmCA2. In contrast, *L. major* malic enzyme (LmME), an unrelated parasite protein (24), was detected only in the corresponding flow-through and not in the immunoprecipitate (Fig. 3G). These findings support a specific association between LmSLC26A and LmCA2 under acidic stress, consistent with their increased colocalization observed by co-immunofluorescence.

### LmSLC26A functions as a bicarbonate transporter and is crucial for maintaining intracellular pH in *Leishmania*

To experimentally verify whether LmSLC26A is indeed the bicarbonate transporter in *L. major*, we planned to knockout the corresponding gene located on chromosome 28 (8). CRISPR/Cas9-mediated gene disruption was performed in the background of an engineered *L. major* strain stably expressing SpCas9 and T7 RNA polymerase (LmCas9:T7Pol) previously generated by us (25, 26). The sgRNAs were designed against the 5′ and 3′ UTRs (28 bp upstream of the start codon and 9 bp downstream of the stop codon, respectively) to enable replacement of the cleaved region with puromycin or blasticidin selection cassettes. Despite repeated attempts, a homozygous LmSLC26A knockout strain could not be obtained under double-antibiotic selection (puromycin + blasticidin). Although this observation alone does not provide definitive evidence for its essentiality, it suggests that complete loss of LmSLC26A may compromise parasite viability. Nevertheless, selection with puromycin alone yielded viable parasites, indicating successful disruption of a single allele and generation of the heterozygous strain (LmSLC26A^+/-^) (Fig. 4A). Genomic PCR analysis using gene-flanking primers yielded two bands of sizes 6.5 kb and 2.3 kb in LmSLC26A^+/-^ parasites, whereas, as expected, only a single 6.5 kb band was detected in the wild type or the control strain (LmCas9:T7Pol), confirming replacement of one allele of LmSLC26A with the puromycin resistance cassette (S5A and B). LmSLC26A transcript and protein levels were reduced by ∼50% in the LmSLC26A^+/-^ strain compared to the control LmCas9:T7Pol cells, as determined by RT PCR and immunofluorescence analysis (Fig. S5C and D, Fig. 4B and C). Given that LmSLC26A contains a predicted histidine phosphatase domain facing the extracellular domain, we checked if membrane phosphatase activity is reduced in the LmSLC26A^+/-^ strain. To specifically assess membrane-associated phosphatase activity, we quantified p-nitrophenyl phosphate (pNPP) hydrolysis using intact cells, thereby restricting the measurement to surface-accessible enzymes. Under this condition, LmSLC26A^+/-^ strain displayed ∼23% reduction in pNPP hydrolytic activity relative to control cells, indicating that LmSLC26A histidine phosphatase domain is functional and contributes substantially to total surface phosphatase activity of the parasite (Fig. 4D). Having confirmed successful generation of the heterozygous strain, we next assessed its bicarbonate uptake capacity using isotope ratio mass spectrometry. While, as expected, basal δ^13^C values were comparable between control and LmSLC26A^+/-^ parasites (Fig S6), following 10 min incubation with ^13^C-bicarbonate, the LmSLC26A^+/-^ strain exhibited a ∼47% reduction in bicarbonate uptake in the LmSLC26A^+/-^ strain with respect to the control cells (Fig. 4E). As reduced bicarbonate uptake is expected to acidify the intracellular environment, we measured intracellular pH in the control and heterozygous strains using BCECF-AM dye. The LmSLC26A^+/-^ strain showed an intracellular pH (pHi) of 6.71 with a calculated proton concentration of 194.99 nmoles/L, whereas the control cells exhibited a near neutral pHi of 6.86 with a corresponding proton concentration of 138.04 nmoles/L. Importantly, the intracellular acidosis observed in the LmSLC26A^+/-^ strain was largely prevented by exogenous bicarbonate, which restored the pHi to 6.82 and reduced the proton concentration to 151.36 nmoles/L (Table 1 and Fig. 4F). Together, these results provide evidence that LmSLC26A functions as a bicarbonate transporter in *Leishmania* and plays a key role in maintaining intracellular pH homeostasis.

**Figure 4.**
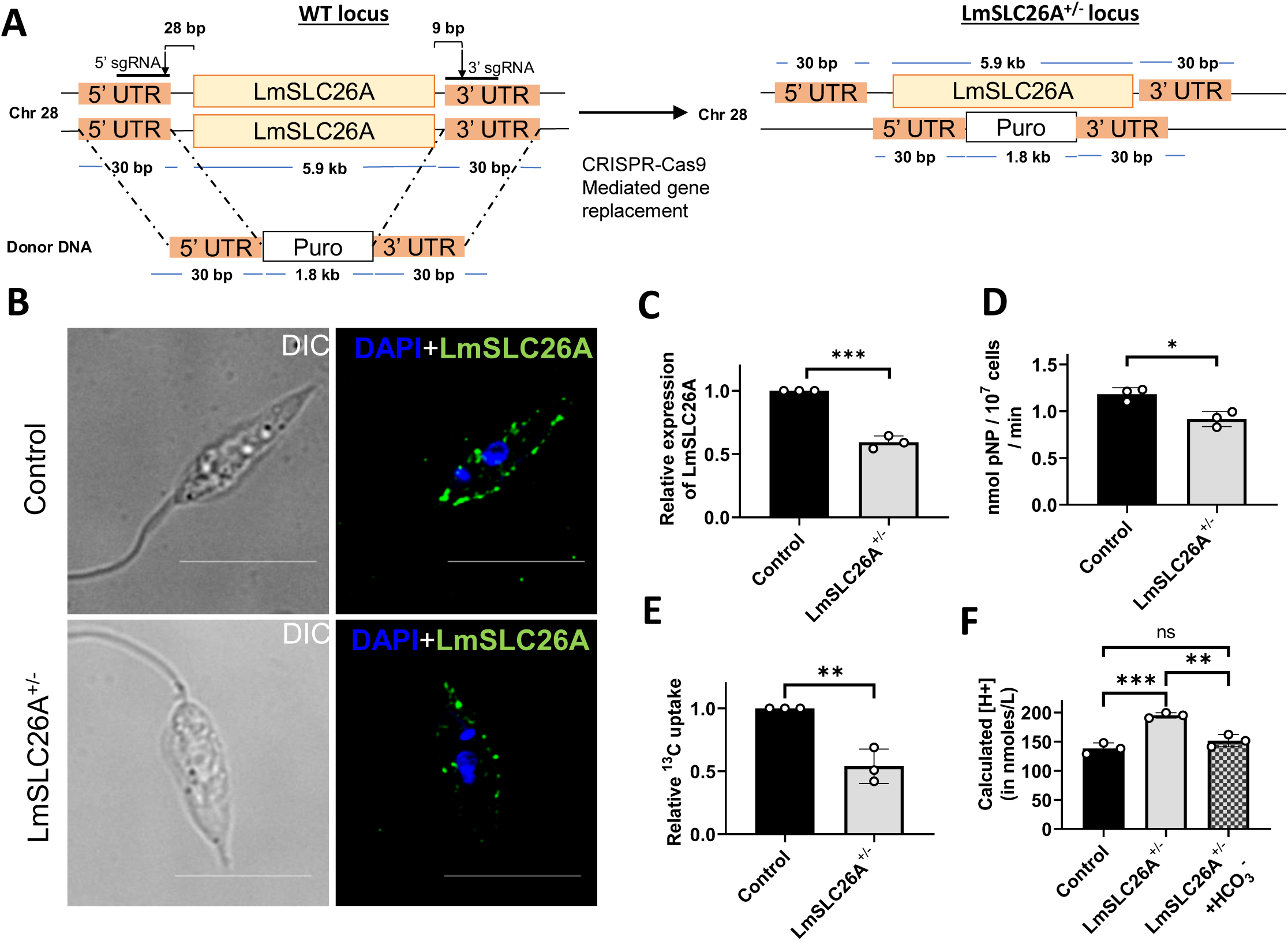
Genetic disruption of LmSLC26A impairs bicarbonate uptake and intracellular pH homeostasis. (A) Schematic illustrating the generation of LmSLC26A^+/-^ cells. Donor DNA containing the puromycin resistance gene flanked by 30 bp homologous regions of LmSLC26A was electroporated into control (LmCas9:T7Pol) cells. Replacement of one allele of LmSLC26A with the puromycin resistance cassette generated the LmSLC26A^+/-^ strain. (B) Control (LmCas9:T7Pol) (top row) and LmSLC26A^+/-^ (bottom row) promastigotes were immunostained with anti-LmSLC26A (green) without permeabilization. DAPI (blue) marks the nucleus and kinetoplast. DIC images (left) are shown alongside corresponding immunofluorescence images (right). Cells were imaged using a Leica SP8 confocal microscope. Scale bar: 10 μm. (C) Quantification of relative LmSLC26A expression based on immunofluorescence intensity. At least 60 cells were analyzed per condition across three independent experiments. Error bars represent SD. ***P = 0.0002. (D) Extracellular p-nitrophenyl phosphate (pNPP) hydrolysis measured in intact promastigotes from control and LmSLC26A^+/-^ strains. Enzyme activity is expressed as nmol pNP per 10^7^ cells. Error bars represent SD from three independent experiments. *P = 0.0135. (E) Comparison of bicarbonate uptake between control (LmCas9:T7Pol) and LmSLC26A^+/-^ strains. Uptake was measured by IRMS following incubation in H^13^CO_3_^-^-containing PBS for 10 minutes. ^13^C uptake was calculated for each strain by subtracting the basal δ^13^C values from δ^13^C values obtained after 10 minutes incubation with ^13^C-bicarbonate. The ^13^C uptake data in LmSLC26A^+/-^ strain is reported relative to the uptake in the control (LmCas9:T7Pol) strain. Error bars represent SD from three independent experiments. **P = 0.0043. (F) Intracellular proton concentration in control (LmCas9:T7Pol), LmSLC26A^+/-^, and LmSLC26A^+/-^ cells grown in bicarbonate-supplemented medium (+HCO_3_^-^), calculated from intracellular pH measurements using BCECF-AM dye. Error bars represent SD from three independent experiments. ***P= 0.0007 (between control and LmSLC26A^+/-^) and **P = 0.0027 (between LmSLC26A^+/-^ and LmSLC26A^+/-^ + HCO3^-^). Statistical significance in all the experiments were determined using Student’s t-test.

**Table 1.** Intracellular pH of different L. major strains^#^.

| ***L. major*** **Strain** | **Measured intracellular pH** |
|---|---|
| Control | 6.86+/- 0.02 |
| LmSLC26A<sup>+/-</sup> | 6.71+/- 0.01** |
| LmSLC26A<sup>+/-</sup> + HCO<sub>3</sub><sup>-</sup> | 6.82+/-0.03 |

<sup>#</sup>Intracellular pH of different *L. major* strains measured using BCECF-AM dye grown for 72 hours in the presence or absence of 5.95 mM bicarbonate. Values represent mean ± SD from three independent experiments. \*\*P = 0.0011(between control and LmSLC26A<sup>+/-</sup>). Statistical significance in all the experiments was determined using Student’s t-test.

### LmSLC26A depletion triggered *Leishmania* cell death and severely impaired parasite virulence *in vitro* and *in vivo*

Having established that LmSLC26A mediates bicarbonate entry into *Leishmania* cells, we next evaluated the physiological consequences of its depletion. The LmSLC26A^+/-^ strain exhibited sluggish growth kinetics, showing ∼30 percent reduced growth relative to control cells at 72 hours. (Fig 5A). Notably, at the same time point, LmSLC26A^+/-^ cell viability was reduced by more than 20%, which correlated with a marked increase in apoptosis in these cells (∼17%) compared to control parasites (∼8%) (Fig. 5B-D). Importantly, supplementation with extracellular bicarbonate substantially ameliorated these defects, restoring growth of the LmSLC26A^+/-^ strain to ∼ 92% of the control level at 72 hours (Fig. 5A). Bicarbonate supplementation also improved viability of the LmSLC26A^+/−^ strain, restoring it to ∼87% of the control level (Fig. 5B), while reducing the proportion of TUNEL-positive cells to ∼13% (Fig. 5C and D). Together, these findings suggest that increased extracellular bicarbonate availability can partially alleviate the physiological consequences of LmSLC26A depletion. These findings are consistent with earlier reports demonstrating intracellular acidosis, induced by carbonic anhydrase inhibition, triggers apoptotic cell death in diverse cancer cell types as well as in *Leishmania* (6, 27–29). Bicarbonate is a well-established activator of soluble adenylyl cyclase, which catalyzes the conversion of ATP to cAMP (30, 31). Given the central role of cAMP signaling in parasite survival and virulence, we examined cAMP levels in the LmSLC26A^+/-^ strain, in which intracellular bicarbonate levels are found to be reduced (32, 33). The LmSLC26A^+/-^ strain displayed ∼60% reduction in intracellular cAMP levels, together with a moderate decrease in ATP levels (Fig. 5E and F). This raises the possibility that impaired bicarbonate transport in the LmSLC26A^+/-^ strain contributes to dampened cAMP signaling and compromised pathogenicity. Furthermore, depletion of LmSLC26A resulted in decreased release of exosomal vesicles, as determined by nanoparticle tracking analysis (NTA) (Fig. 5G). This observation is intriguing, as *Leishmania* is known to secrete numerous virulence factors via exosome mediated pathways, suggesting that reduced vesicle release may contribute to the diminished virulence of the LmSLC26A^+/-^ parasite (34, 35).

**Figure 5.**
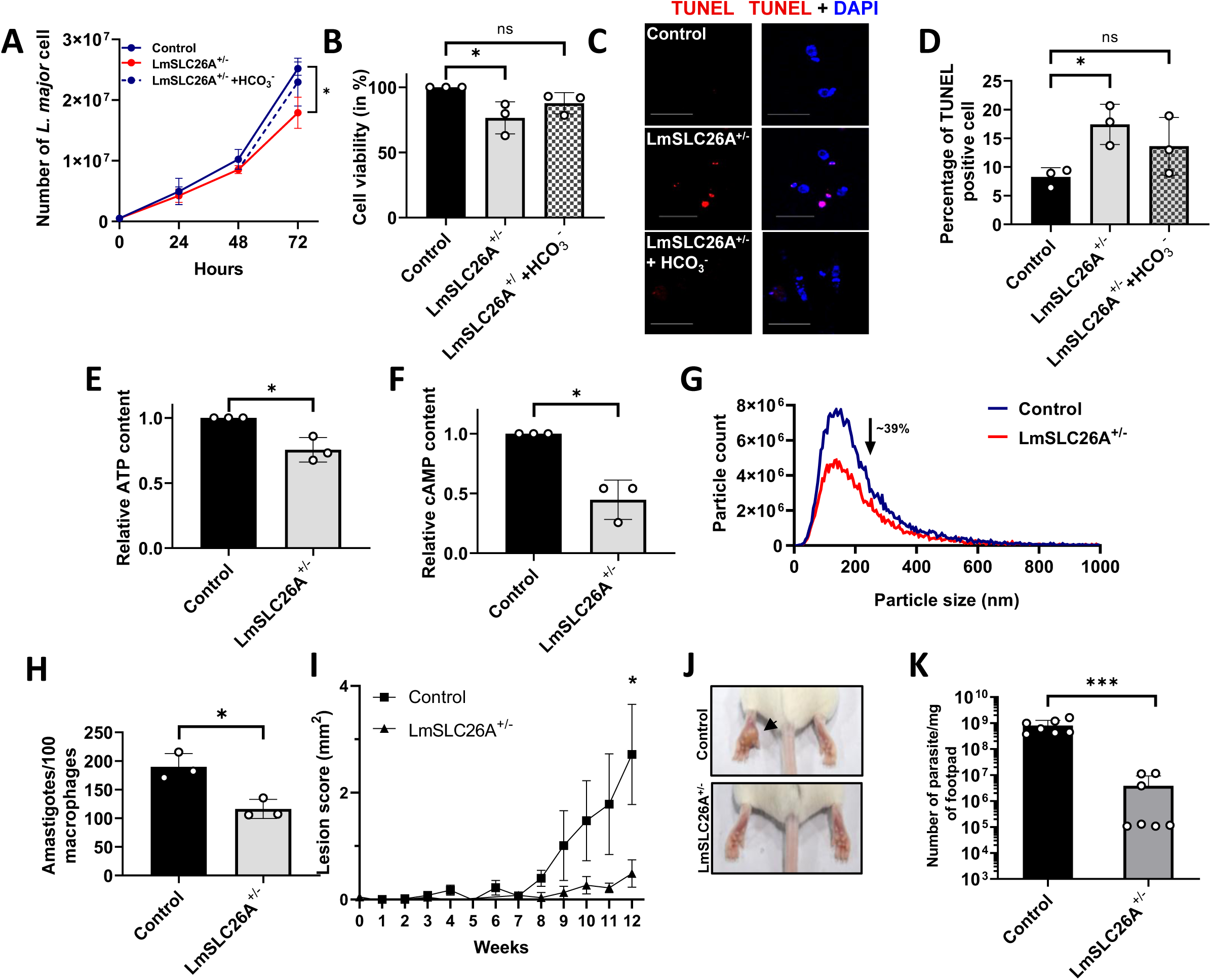
LmSLC26A is critical for parasite survival, metabolic fitness, and virulence. (A) Control (LmCas9:T7Pol), LmSLC26A^+/-^ and LmSLC26A^+/-^ promastigotes supplemented with 5.95 mM bicarbonate were cultured for 72 h, and cell numbers were quantified every 24 h. Error bars represent SD from three independent experiments. *P = 0.01077. (B) Cell viability was assessed by MTT assay after 72 h. Error bars represent SD from three independent experiments. *P = 0.0304, ns, P = 0.0607. (C) Representative TUNEL assay images of control (LmCas9:T7Pol) (top row), LmSLC26A^+/-^ (middle row) and LmSLC26A^+/-^ promastigotes supplemented with 5.95 mM bicarbonate (bottom row). TUNEL staining (red) and merged images with DAPI (blue) are shown. (D) Quantification of TUNEL-positive cells reveals an increased proportion of apoptotic cells in the LmSLC26A^+/-^ strain. Error bars represent SD from three independent experiments. *P = 0.0146, ns, P = 0.1532. (E) Relative intracellular ATP levels in control (LmCas9:T7Pol) and LmSLC26A^+/-^ strains. Error bars represent SD from three independent experiments. *P = 0.0109. (F) Relative intracellular cAMP levels in control (LmCas9:T7Pol) and LmSLC26A^+/-^ strains. Error bars represent SD from three independent experiments. *P = 0.0044. (G) Representative size distribution profiles of extracellular vesicles (EVs) released by control (LmCas9:T7Pol) and LmSLC26A^+/-^ strains, measured by nanoparticle tracking analysis using the NanoSight Pro. The LmSLC26A^+/-^ strain exhibits a ∼39% reduction in particle counts within the 50-200 nm range. Profiles represent averages from three independent biological replicates. (H) Intracellular parasite burden in J774A.1 macrophages infected with control (LmCas9:T7Pol) or LmSLC26A^+/-^ strains. Cells were fixed and stained with DAPI, and the number of intracellular amastigotes per 100 macrophages was determined from epifluorescence images. Error bars represent SD from three independent experiments. *P = 0.0114. (I) *In vivo* infectivity assay: control (LmCas9:T7Pol) or LmSLC26A^+/-^ parasites were injected into the left hind footpad of BALB/c mice, and lesion progression was monitored for 12 weeks. Lesion scores were recorded weekly. Error bars represent SEM (n = 7 mice per group). P values for weeks 8, 9, 10, and 11 are 0.0754, 0.1747, 0.1353, and 0.1094, respectively. *P = 0.0461 (week 12). (J) Representative footpad images at 12 weeks post-infection for control (top) and LmSLC26A^+/-^ (bottom) groups. The black arrow indicates swelling in the footpad infected with the control strain. (K) Parasite burden in infected footpad tissue at 12 weeks post-infection, determined by serial dilution assay and expressed as parasites per mg of tissue. Data represent mean ± SD (n = 7 mice per group). ***P = 0.0007. Statistical significance in all the experiments were determined using Student’s t-test.

To check the impact of LmSLC26A depletion on parasite virulence, J774A.1 macrophages were infected with control or LmSLC26A^+/-^ *L. major*, and intracellular parasite burdens were quantified. Infection with the LmSLC26A^+/-^ strain resulted in ∼30% reduction in intracellular parasite burden compared with the control, indicating impaired parasite survival within host cells (Fig. 5H). Consistent with this attenuated virulence observed *in vitro*, our *in vivo* infection experiments further revealed drastically reduced pathogenicity of the LmSLC26A^+/-^ strain. While BALB/c mice infected with control *L. major* developed progressive footpad lesions, reaching a mean lesion score of ∼2.7 mm^2^ at 12 weeks post infection, mice infected with the LmSLC26A^+/-^ strain exhibited significantly diminished lesion development, with a mean lesion score of only ∼0.5 mm^2^ at the same time point (Fig. 5I and J). Corroborating the reduced lesion size, parasite burden in footpad tissues infected with the LmSLC26A^+/-^ strain was almost 100-folds lower than in control infections (Fig. 5K). These data validate that diminished lesion development in LmSLC26A^+/-^ infected mice is due to impaired parasite survival rather than attenuated inflammatory responses. This is likely driven by the combined effects of increased parasite cell death, reduced cAMP levels, and diminished exosome-mediated release of virulence factors in the LmSLC26A^+/-^ strain. Together, our *in vitro* and *in vivo* infection experiments establish LmSLC26A mediated bicarbonate uptake as a critical determinant of intracellular persistence of *Leishmania* and its virulence.

## Discussion

The ability of *Leishmania* to survive within the acidic phagolysosomal environment of host macrophages necessitates robust mechanisms to regulate intracellular pH homeostasis. Yet, how the parasite maintain a near neutral cytosolic pH under such extreme conditions remains incompletely understood. While earlier studies primarily attributed cytosolic pH regulation to proton extrusion mediated by plasma membrane H^+^-ATPases, our recent findings supported a bicarbonate-based buffering mechanism, driven by the coordinated action of two carbonic anhydrases, cytosolic LmCA1, and membrane-bound LmCA2 (5, 6). This model predicted the existence of a plasma membrane bicarbonate transporter that would complete the buffering cycle by mediating bicarbonate re-entry into the parasite. However, in the absence of any annotated bicarbonate transporter in the parasite genome the identity of this critical transporter in *Leishmania* has remained elusive. Here, we provide strong evidence that LmSLC26A fulfils this role, thereby defining the molecular basis of the bicarbonate buffering mechanism that enables *Leishmania* to maintain cytosolic pH neutrality and sustain its virulence. LmSLC26A represents the first experimentally validated bicarbonate transporter identified in kinetoplastid and entire protozoan lineages.

The presence of an active bicarbonate transport system in *Leishmania* was supported by our initial findings, demonstrating: (a) treatment with DIDS, a bicarbonate transport inhibitor, resulted in a dose dependent reduction in parasite growth; and (b) *Leishmania* cells could directly uptake ^13^C bicarbonate, confirmed by IRMS-based analysis. Interestingly, extracellular bicarbonate partially restored parasite growth even in the presence of DIDS, suggesting that increased bicarbonate availability could overcome its inhibitory effect. A recent structural study of anion exchanger 1 (AE1/SLC4A1), a well-characterized Cl⁻/HCO₃⁻ exchanger, demonstrated that DIDS occupies the anion-access channel, with one of its sulfonic acid groups positioned within 4 Å of the bound bicarbonate ion (36). It was also shown that DIDS-mediated inhibition of SLC4 transporters is rapid and reversible due to ionic interactions with lysine residues in the anion-access pathway (37). These findings raise the possibility of a competitive component of inhibition, whereby increased extracellular bicarbonate may partially overcome DIDS binding. Although this mechanism has not been directly established for LmSLC26A, it provides a plausible explanation for the observed bicarbonate-mediated rescue. Additionally, residual transporter activity, passive CO_2_ diffusion across plasma membrane followed by intracellular hydration, and alternative bicarbonate uptake pathways may also contribute to the observed rescue. Together, these observations provided a strong conceptual framework for uncovering the molecular identity of the bicarbonate transporter in *Leishmania*.

Despite its annotation as a “sulfate transporter-like protein” in the *L. major* genome, LmSLC26A showed strong sequence homology to human SLC26 family members, which function as versatile multi-anion exchangers (10). LmSLC26A also retains hallmark structural features of this family, including multiple transmembrane domains and a conserved STAS domain. Building on these bioinformatics analyses, our imaging data demonstrated that LmSLC26A is localized to the parasite plasma membrane, positioning it to directly mediate bicarbonate influx from the extracellular environment. Consistent with this localization, ^13^C-bicarbonate uptake into *Leishmania* cells was significantly impaired upon genetic depletion of LmSLC26A, leading to intracellular acidification and thereby establishing its central role in cytosolic pH homeostasis. Importantly, supplementation with exogenous bicarbonate restored intracellular pH in the mutant parasites, indicating that increased extracellular bicarbonate availability can compensate for the defect associated with reduced LmSLC26A expression. This rescue could be mediated, at least in part, by the remaining functional LmSLC26A allele. However, contributions from additional bicarbonate transport mechanisms or CO_2_ diffusion followed by cytosolic LmCA1-mediated hydration cannot be excluded. Thus, although these findings support an important role for LmSLC26A in bicarbonate acquisition by the parasite, they do not establish it as the sole route of bicarbonate entry. A relevant precedent exists in cyanobacteria, where the bicarbonate transporter BicA was initially misannotated as a sulfate transporter based on sequence similarity but was subsequently redefined through functional characterization (14). Interestingly, under acidic conditions, LmSLC26A was found to associate with the membrane-bound carbonic anhydrase LmCA2, suggesting the formation of a functional transport complex analogous to bicarbonate transport metabolons described in mammalian systems (22, 23). This interaction likely supports an integrated mechanism for efficient cytosolic buffering, enabling parasite adaptation within the hostile environment of the host phagolysosome. The elevated levels of LmSLC26A under acidic conditions further suggest that its role may be particularly important in the amastigotes, although the mechanisms governing its pH-dependent regulation remain unclear.

Depletion of LmSLC26A had pleiotropic effects on parasite physiology beyond disrupting pH homeostasis. LmSLC26A-deficient parasites exhibited a moderate reduction in growth. Given that bicarbonate serves not only as a buffering agent but also as a substrate for key carboxylation reactions, its impaired uptake is likely to perturb central metabolic pathways, contributing to the observed growth defect (38). Alternatively, this effect may arise indirectly from intracellular pH imbalance, which can subtly alter metabolic reactions (39). In this context, it is worth noting that an ongoing genome-wide bar-seq deletion study by the LeishGEM team also reports a fitness defect in *L. mexicana* promastigotes lacking the orthologous gene (LmxM.28.1690), as evidenced by a progressive decline in representation of the mutant strain within the pooled population (40). LmSLC26A depleted parasites also displayed features of apoptosis, suggesting that intracellular acidification and the resulting metabolic imbalance act as critical triggers of cell death. This is consistent with multiple reports demonstrating that disruption of carbonic anhydrase-mediated pH regulation led to intracellular acidification and inducing apoptosis in *Leishmania* as well as in cancer cells (6, 27–29). Interestingly, LmSLC26A-depleted parasites also exhibited moderately reduced intracellular ATP levels, which may reflect secondary metabolic consequences of impaired pH homeostasis, as intracellular pH is closely linked to cellular energy metabolism and mitochondrial bioenergetics (41).

One of the most striking phenotypes observed in the LmSLC26A^+/-^ strain was a marked reduction in intracellular cAMP levels. Bicarbonate is known to activate soluble adenylyl cyclases in diverse organisms, thereby promoting the conversion of ATP to cAMP (30, 31). In *L. major*, the presence of a heme-containing soluble adenylyl cyclase (HemAC-Lm) raises the possibility that intracellular bicarbonate may influence cAMP production (33). Thus, reduced bicarbonate availability following LmSLC26A depletion could potentially affect HemAC-Lm activity and contribute to the observed decrease in intracellular cAMP. Given the central role of cAMP in regulating oxidative stress tolerance, stage differentiation, intracellular survival, and infectivity in *Leishmania*, altered cAMP homeostasis may contribute to the impaired parasite physiology and attenuated virulence observed in the LmSLC26A^+/-^ strain (32, 42). We also observed reduced exosomal vesicle release in the LmSLC26A^+/-^ strain, although the underlying basis for this intriguing phenotype remains unclear. As *Leishmania* secretes a large number of virulence factors via exosome-mediated pathways (34, 35), reduced vesicle release may represent an additional factor potentially contributing to the attenuated pathogenicity of the LmSLC26A^+/-^ parasite.

Consistent with these findings, LmSLC26A-depleted parasites exhibited reduced survival within macrophages and markedly attenuated virulence in mice, as evidenced by diminished lesion development and a substantial reduction in parasite burden in the infected tissue. Notably, this pronounced loss of pathogenicity arise despite only a ∼50% reduction in LmSLC26A levels in the mutant strain and likely reflect the combined effects of increased cell death, decreased cAMP levels, and reduced exosomal release of virulence factors. Bicarbonate has previously been linked to virulence in bacterial pathogens such as *Vibrio cholerae* and *Staphylococcus aureus*, where bicarbonate sensing modulates virulence gene expression (43, 44). Our study extends this paradigm to protozoan parasites and identified bicarbonate transport as a key determinant of pathogenicity in *Leishmania*, highlighting it as a potential target for therapeutic intervention.

Overall, our study identifies LmSLC26A as an atypical and multifunctional member of the SLC26 family that warrants further investigation. Unlike canonical counterparts, LmSLC26A is considerably larger in size and harbours a histidine ecto-phosphatase domain, suggesting functional diversification beyond classical anion transport. Supporting this, the LmSLC26A^+/-^ strain exhibited almost 23% reduction in surface phosphatase activity, demonstrating that this domain is enzymatically active and contributes substantially to total phosphatase activity at the parasite membrane. In *Leishmania*, ecto-phosphatases are known to facilitate phosphate acquisition from the host environment and to participate in host-parasite interactions, including macrophage invasion and intracellular survival (45, 46). The fusion of such an enzymatic domain with a bicarbonate transporter is unprecedented and points to a previously unrecognized class of multifunctional membrane proteins. These observations raise the possibility that LmSLC26A integrates pH regulation, nutrient acquisition, and enzymatic activity within a single molecular framework. Accordingly, some of the phenotypic consequences of LmSLC26A depletion may also arise from reduced phosphatase activity in addition to impaired bicarbonate transport. Although our data demonstrate reduced bicarbonate uptake and intracellular acidification following LmSLC26A depletion, the relative contributions of these two activities to the observed phenotypes, particularly parasite virulence, remain to be experimentally determined. Moreover, the transport kinetics, substrate specificity, and dependence of bicarbonate transport on energy or ion gradients remain to be characterized. Future studies involving mutational analysis of conserved residues implicated in bicarbonate transport and functional dissection of the histidine phosphatase domain will be important to delineate the mechanistic contributions of these two activities and further elucidate how LmSLC26A coordinates multiple aspects of parasite physiology and virulence.

### Experimental procedures

Unless otherwise mentioned, all reagents were purchased from Sigma-Aldrich. Details of all primers and antibodies used in this study are provided in Tables S2 and S3, respectively.

### Ethics statement

The *L. major* infection studies in BALB/c mice and antibody generation in mice and rabbit were approved by the Institutional Animal Ethics Committee (IAEC) and conducted in accordance with Committee for Control and Supervision of Experiments on Animals (CCSEA) guidelines.

### Cell culture, growth kinetics and cell viability assay

*L. major* (MHOM/SU/73/5-ASKH) cells, J774A.1 murine macrophages and HEK-293 cells were obtained from ATCC. As, described previously *L. major* promastigotes were cultured at 26°C in M199 medium supplemented with 15% heat-inactivated fetal bovine serum, 23.5 mM HEPES, 0.2 mM adenine, 150 µg/ml folic acid, 10 µg/ml hemin, 120 U/ml penicillin, 120 µg/ml streptomycin, and 60 µg/ml gentamicin (29). For growing cells under acidic condition, the pH of the medium was adjusted to 5.5 using diluted HCl. Wherever indicated, sodium bicarbonate (5.95 mM) was added to the medium and the pH was readjusted to either 7.2 or 5.5.

DIDS was freshly prepared in DMSO and diluted in water, as required. *L. major* promastigotes were incubated with DIDS containing medium at the indicated concentrations for 72 h, after which cell growth was assessed using a hemocytometer. Control cultures received an equivalent concentration of DMSO.

*L. major* promastigotes were seeded at a density of 5 × 10^5^ cells/ml and cell numbers were determined at 24, 48, and 72 h using a hemocytometer after fixation in 0.9% formal saline. Cell viability was assessed by the MTT assay, as described previously (6). Briefly, 4 × 10^6^ log phase cells from each strain were incubated with 0.5 mg/ml MTT for 3 h. Cells were then harvested, washed with PBS, and resuspended in 0.04 N HCl in isopropanol to solubilize the formazan product. Absorbance was measured at 595 nm using a microplate reader. Percentage viability of mutant strains was calculated relative to the control strain (OD-mutant/OD-control × 100).

J774A.1 murine macrophages were cultured in Dulbecco’s modified Eagle’s medium (DMEM) supplemented with 10% heat-inactivated fetal bovine serum, 2 mM L-glutamine, 0.91 mM sodium pyruvate, 100 μg/ml streptomycin, and 100 U/ml penicillin. HEK293 cells were cultured in Minimum essential medium (MEM) supplemented with 10% heat-inactivated fetal bovine serum, 2 mM L-glutamine, 0.91 mM sodium pyruvate, 100 μg/ml streptomycin, and 100 U/ml penicillin. Both cell lines were maintained at 37°C in a humidified atmosphere containing 5% CO_2_.

### 13C-bicarbonate uptake assay by isotope ratio mass spectrometry

Given that ^12^C is the predominant naturally occurring carbon isotope (∼98.9%), we used ^13^C-bicarbonate (98 atom % ^13^C, 99% purity) as a tracer and developed an isotope ratio mass spectrometry (IRMS)-based bicarbonate uptake assay. Briefly, 2 × 10^7^ *L. major* promastigotes were harvested and washed with PBS. Cells were incubated in 5.95 mM ^13^C-bicarbonate for 2.5, 5 and 10 min, followed by centrifugation at 1,000 × g for 5 min at 4 °C. The cell pellet was washed with PBS and centrifuged again. Cells were then resuspended in 10 µl of PBS, and the concentrated suspension was distributed into three tin capsules, tightly sealed, and immediately loaded into an organic elemental analyzer (Flash 2000, Thermo Fisher Scientific). The carbon isotopic value (δ^13^C) of the resulting samples were subsequently analyzed in a Delta V Plus isotope ratio mass spectrometer (IRMS) (Thermo Fisher Scientific) coupled to the elemental analyzer via Conflo IV device (Thermo Fisher Scientific). For measuring the basal δ^13^C value, *Leishmania* promastigotes were directly analyzed in IRMS without addition of ^13^C-bicarbonate. The relative abundance of the stable carbon isotope ^13^C compared to ^12^C in sample to that of the standard were expressed as δ^13^C values, reported in per mille (‰) against the Vienna Pee Dee Belemnite (VPDB) international reference scale using the equation (15):

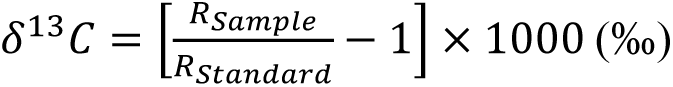

where, *R_Sample_* = (^13^C/^12^C) ratio of sample and *R_Standard_* is (^13^C/^12^C) ratio of the standard. δ^13^C values are reported in per mille (‰), which enables standardized comparison of isotopic measurements across experiments and laboratories.

During the entire run, international standards of known δ^13^C values were analyzed after a batch of five samples to ensure accurate measurement of δ^13^C values. International Atomic Energy Agency (IAEA) standards CH3 (δ^13^C=-24.72 ‰ VPDB), CH6 (δ^13^C=-10.449 ‰ VPDB) and Caffeine (δ^13^C=-27.771 ‰ VPDB) were used for this study. The instrument precision was ∼0.2 ‰.

### RNA isolation and RT-PCR

Total RNA from *L. major* promastigotes was isolated using TRIzol reagent (Invitrogen) and treated with DNase I (Invitrogen). cDNA was synthesized from DNase I-treated RNA using the Verso cDNA kit (Thermo Scientific). Minus-reverse transcriptase (-RT) controls were prepared by omitting reverse transcriptase while including all other reaction components. PCR was performed using cDNA derived from both +RT and -RT samples using Leo Master Mix (Dxbidt) for 18-24 cycles. LmSLC26A transcripts were amplified using internal primers P7/P8, whereas rRNA45 was amplified using primers P9/P10.

### Bioinformatics studies

Sequence alignments were performed using Clustal Omega and visualized in Inkscape. Protein sequences were retrieved from UniProt (47). Mutated residues in human SLC26A3 were compiled from the public version of the Human Gene Mutation Database (48) and mapped onto both the sequence alignment and the domain architecture. Transmembrane regions and the STAS domain were annotated based on UniProt, while the histidine phosphatase domain was defined using InterPro. For analysis of the histidine phosphatase domain, the region corresponding to residues 730 to 999 of LmSLC26A was extracted and aligned with homologous domains from characterized bacterial and eukaryotic histidine phosphatases.

### Antibody generation

Polyclonal antibodies were generated against fragments of LmSLC26A (amino acids 597-935) and LmCA2 (amino acids 291-540). The corresponding regions were PCR-amplified using primers P3/P4 (for cloning of LmSLC26A^597-935^) or P5/P6 (for cloning of LmCA2^291-540^) from *L. major* genomic DNA and cloned into the pET28a expression vector. Recombinant constructs were transformed into *E. coli* BL21 (DE3) cells, and protein expression was induced with 0.5 mM IPTG at 20 °C for 12 h. Cells were harvested, lysed, and the recombinant protein fragments were purified by Ni-NTA affinity chromatography following standard procedures. For LmSLC26A^597-935^, purification was carried out under denaturing conditions with 8 M urea included in the lysis, wash, and elution buffers to enhance solubility and yield. Urea was subsequently removed by dialysis. Protein concentrations were determined by the Lowry method, and purity was assessed by SDS-PAGE.

For immunization, 4-6-week-old female BALB/c mice were injected subcutaneously with 50 µg of recombinant protein (LmSLC26A^597-935^) emulsified in complete Freund’s adjuvant (Santa Cruz Biotechnology). Three booster doses were administered at two-week intervals using incomplete Freund’s adjuvant. For rabbit immunization, 6-12-month-old animals were injected with 100 µg of recombinant proteins (LmSLC26A^597-935^ or LmCA2^291-540^), prepared similarly with complete Freund’s adjuvant, followed by three booster injections at two-week intervals with incomplete Freund’s adjuvant. Serum was collected 7 days after the final injection and stored at -80 °C until further use.

### Generation of HEK-293 cells stably expressing LmSLC26A

LmSLC26A^1–1024^ was PCR amplified from *L. major* genomic DNA and was cloned into the EcoRI site of the pCXN vector using primers P1/P11. The recombinant construct was verified by restriction digestion and used for transfection into HEK-293 cells. For transfection, 1 × 10⁶ cells were seeded in 6-well plates containing MEM supplemented with 2% FBS without antibiotics and incubated overnight until the cells reached approximately 70–80% confluency. The medium was replaced with fresh MEM (1.7–1.8 ml per well) before transfection. Plasmid DNA (2 µg) was diluted in 100–150 µl of MEM, while Lipofectamine 2000 (2 µl) was diluted separately in an equal volume of MEM. The diluted DNA and Lipofectamine 2000 solutions were combined, mixed gently, and incubated at room temperature for 20 min to allow DNA-lipid complex formation. The complexes were then added dropwise to the cells, followed by gentle swirling of the plates to ensure even distribution. Cells were incubated at 37 °C in a humidified atmosphere containing 5% CO₂ for 48 h. Following incubation, the medium was replaced with fresh MEM containing 600 µg/ml G418 sulphate for selection of stably transfected cells. The selection medium was replaced every 3 days until resistant colonies appeared. Individual colonies were isolated using cloning discs and transferred to separate 35-mm culture dishes for expansion, followed by further expansion in 60-mm culture dishes. A fraction of the cells from each clone was used to assess LmSLC26A expression by immunofluorescence using anti-LmSLC26A antibodies.

### Immunofluorescence microscopy

*L. major* promastigotes were allowed to adhere to poly-L-lysine-coated coverslips for ∼30 min at 26 °C, fixed with 4% paraformaldehyde for 10 min in the dark, washed with PBS, and blocked with 0.2% gelatin for 10 min. Cells were incubated with primary antibodies for 90 min, washed, and then incubated with appropriate secondary antibodies for 90 min. After washing, coverslips were mounted in antifade medium containing DAPI (Vectashield). Images were acquired using a Leica SP8 confocal microscope with a 63× oil-immersion objective. Image processing and analysis were performed using LAS X software or ImageJ. For each sample, 1-3 central z-stacks were merged, and maximum-intensity projections were generated for presentation. For amastigotes, a similar procedure was followed. Parasites were isolated from infected footpad tissue by mechanical disruption, resuspended in PBS, and allowed to adhere to coverslips prior to fixation with 4% paraformaldehyde. Prepared coverslips were stored at 4 °C until further use. Colocalization analysis was performed in ImageJ using the JaCoP plugin.

### Coimmunoprecipitation and western blotting

For coimmunoprecipitation, 2 x 10^8^ *L. major* promastigotes were harvested, washed with PBS, and resuspended in 1 ml of coimmunoprecipitation lysis buffer containing 50 mM Tris (pH 7.4), 150 mM NaCl, 1 mM EDTA, 1% NP-40, 1% sodium deoxycholate, and 0.1% SDS, with 1X protease inhibitor cocktail). The lysate was gently sonicated three times and centrifuged at 14,000 × g for 30 min at 4 °C to remove insoluble debris. The supernatant was collected, and 10 µl of primary antibody serum was added to 300 µl of the clarified lysate, followed by overnight incubation at 4 °C with gentle rotation. The following day, Protein A-agarose beads (BioBharati) were added and incubated for 2 h at 4 °C with rotation. The beads were then pelleted by centrifugation at 1,000 × g for 4 min at 4 °C and washed 2-3 times with lysis buffer. Bound proteins were eluted using 0.1 M glycine (pH 2.0), and the eluted fractions were subsequently analyzed by western blotting.

Protein fractions were resolved by SDS-PAGE, transferred to PVDF membranes, and blocked with 5% skimmed milk in TBST, followed by incubation with primary antibodies overnight at 4 °C. Following washes, membranes were incubated with HRP-conjugated secondary antibodies for 90 min at room temperature. Signals were detected using a chemiluminescent substrate (Bio-Rad) and imaged on a ChemiDoc system (Bio-Rad).

### Generation of LmSLC26A^+/-^ heterozygous *L. major* strain

The LmSLC26A^+/-^ heterozygous strain was generated in the background of a *L. major* strain stably expressing Cas9 and T7 RNA polymerase (LmCas9:T7Pol), previously developed by our group, following the method described by Beneke et al. (25, 26). Primers for amplification of sgRNA templates were designed using the LeishGEdit primer design module, and the sequences are provided in Table S2. Briefly, 5′ and 3′ sgRNA oligonucleotides were PCR-amplified using primer G00 with P12 or P13 to generate sgRNA templates. In parallel, puromycin- and blasticidin-resistance cassettes, each containing 30-nt overhangs corresponding to the 5′ and 3′ UTRs of LmSLC26A, were PCR-amplified from pTPuro and pTBlast plasmids using primers P14 and P15. All four PCR products, the 5′ sgRNA template, 3′ sgRNA template, and the two resistance cassettes, were co-electroporated into log-phase LmCas9:T7Pol promastigotes using a standard protocol. The strain was selected in presence of Blasticidin (5 µg/ml), puromycin (20 µg/ml), or both. The LmSLC26A^+/-^ heterozygous strain was verified at the genomic, RNA, and protein levels, as detailed in the results section.

### Determination of *L. major* surface phosphatase activity

8 × 10^6^ *L. major* promastigotes were harvested by centrifugation at 1,000 × g for 5 min and washed three times with wash buffer containing 20 mM HEPES and 150 mM NaCl (pH 7.2). Cells were then gently resuspended in assay buffer containing 50 mM sodium acetate and 150 mM NaCl (pH 5.5). The reaction was initiated by addition of p-nitrophenyl phosphate (pNPP; Amresco) to a final concentration of 5 mM, followed by incubation at 26 °C for 30 min. Cells were subsequently removed by centrifugation, and the supernatant was transferred to a 96-well plate. The reaction was terminated by addition of 1 M NaOH, which simultaneously converted p-nitrophenol to its deprotonated form. Absorbance was measured at 405 nm using a spectrophotometer. Enzymatic activity was calculated as p-nitrophenol equivalents using a standard curve generated with known concentrations of p-nitrophenol (HiMedia) prepared in the same buffer.

### Intracellular pH measurement

Intracellular pH of *L. major* promastigotes was measured using the fluorescent probe BCECF-AM (2′,7′-bis-(2-carboxyethyl)-5-(and 6)-carboxyfluorescein acetoxymethyl ester) (Invitrogen) as described previously (6). Briefly, 1 × 10^7^ promastigotes were washed with PBS and resuspended in 500 μl of buffer A (136 mM NaCl, 2.68 mM KCl, 0.8 mM MgSO_4_, 11.1 mM glucose, 1.47 mM KH_2_PO_4_, 8.46 mM Na_2_HPO_4_, 1 mM CaCl_2_, and 20 mM HEPES, pH 7.0). Cells were incubated with 10 μM BCECF-AM for 30 min at 26 °C, washed twice with buffer A, and resuspended in 100 μl of the same buffer. Fluorescence was measured using a BioTek Cytation 5 microplate reader with excitation at 490 nm (pH sensitive) and 440 nm (isosbestic point), and emission at 535 nm. The ratio of fluorescence intensities (490/440) was converted to intracellular pH using a calibration curve. For calibration, BCECF-loaded cells were incubated in potassium phosphate buffers of defined pH (6.5 to 7.5) in the presence of 5 μg/ml nigericin, and fluorescence ratios were recorded as a function of pH.

### TUNEL assay

The TUNEL assay to detect apoptotic cells was performed using the In Situ Cell Death Detection Kit, TMR Red (Roche). Briefly, *L. major* cells were fixed with 4% paraformaldehyde for 10 minutes, washed twice with PBS, and permeabilized with 0.1% Triton X-100 for 2 minutes, followed by an additional PBS wash. Freshly prepared TUNEL reaction mixture was then added and incubated for 60 minutes at 37°C. Thereafter, cells were washed twice with PBS, spread onto poly-L-lysine-coated coverslips, and mounted using Vectashield containing DAPI. Imaging was performed using either a Leica confocal microscope or an IX81 epifluorescence microscope, and only nuclear-localized red fluorescence signals were considered TUNEL-positive.

### Intracellular cAMP and ATP measurement

1 × 10^7^ *L. major* promastigotes were lysed by the freeze-thaw method, and whole-cell lysates were used for cAMP quantification using a commercial kit (Elabscience®) according to the manufacturer’s instructions. cAMP levels were normalized to total protein content determined by the Folin-Lowry method.

Intracellular ATP levels in *L. major* promastigotes were measured using an ATP determination kit (Invitrogen) as described previously (24). Briefly, 4 × 10^7^ cells were resuspended in 50 µl of PBS and lysed by sonication. 10 µl lysate was added to freshly prepared ATP reaction solution and incubated for 15 min at room temperature. Luminescence was measured at 560 nm, and ATP concentrations were calculated from a standard calibration curve.

### Nanoparticle tracking analysis of *Leishmania* culture supernatant

*L. major* promastigotes were seeded at a density of 5 × 10^6^ cells/ml and cultured for 72 hours. A total of 1 × 10^7^ cells were then harvested, washed with particle-depleted medium (prepared by ultracentrifugation of M199 medium at 100,000 × g for 1 hour, followed by collection of the supernatant and sterile filtration), and finally resuspended in 1 ml of the same medium. Cells were incubated at 26 °C for another 12 hours, after which they were counted to confirm cell density and centrifuged at 1500 × g for 5 minutes at 4 °C. The supernatant medium was carefully collected and subjected to nanoparticle tracking analysis using the NanoSight Pro. Prior to data acquisition, the flow cell was equilibrated with the respective supernatant. For each sample, 10 videos were recorded in light scatter mode at a flow rate of 3 µl/min. Between captures, samples were advanced at 20 µl/min for 8 seconds followed by a 3 second stabilization period. Focus and exposure settings were optimized and maintained consistently across all samples within an experiment. Particle size distribution and concentration were analyzed using the instrument software and plotted using GraphPad Prism.

### *Leishmania* infection experiments

*L. major* infection of J774A.1 macrophages was performed as described previously (6). Briefly, macrophages were activated with 100 ng/ml LPS for 6 hours and infected with stationary phase promastigotes at a multiplicity of infection (MOI) of 1:30. After 12 hours, uninternalized parasites were removed by washing with PBS, and cells were incubated in fresh DMEM for ∼24 hours. Cells were then fixed in 1:1 (v/v) acetone:methanol for 10 minutes and mounted with DAPI-containing medium. Infection was quantified by imaging at least 100 macrophage nuclei from multiple fields using an Olympus IX81 epifluorescence microscope and calculating the number of amastigote nuclei per 100 macrophage nuclei from three independent experiments.

For *in vivo* infection studies, 5 × 10^6^ late stationary phase promastigotes were injected into the left hind footpad of 4-6 week old BALB/c mice. Footpad swelling was measured weekly using calipers, and lesion size was calculated as (infected − uninfected thickness) × (infected − uninfected width). At 12 weeks post infection, mice were euthanized and parasite burden was determined by limiting dilution (49). Excised footpads were washed, weighed, homogenized in complete M199, serially diluted, and plated in 96 well plates. Parasite load was estimated from the highest dilution showing growth after 10 days at 26 °C and expressed as parasites per milligram of tissue. Each group contained seven mice.

### Statistical Analysis

Statistical analyses were performed using Microsoft Excel 2019. Comparisons between the two groups in all experiments were conducted using unpaired, two-tailed Student’s t-test. p-values are reported and indicated in the figure legends. Data are presented as mean ± standard deviation (SD).

## Data availability

Data used in the study are included in the manuscript and/or supporting information.

## Supporting information

This article contains supporting information.

## Acknowledgments

We sincerely thank Prof. Prasanta Sanyal, Mr. Mahesh Ghosh, and the IRMS Central Facility, IISER Kolkata, for assistance with the bicarbonate uptake study. We thank Mr. Jajati Keshari Ray, Mr. Anu Das, and the IISER Kolkata Animal Facility for assistance with animal studies. We thank Drs. Subrata Adak, Sankar Maiti, and Subhankar Dolai for providing various reagents. Department of Biotechnology-Builder funded (BT/INF/22/SP45383/2022) IISER Kolkata core imaging facility is also thankfully acknowledged.

## Funding and additional information

This work was supported by funding from DST (EMR/2017/004506), ICMR (6/9-7(318)/2023-ECD-II), and West Bengal DSTBT (398(Sanc.)/STBT-38113015/16/2024-ST SEC) awarded to RD. AS and SPA were supported by CSIR fellowship.

## Conflict of interest

The authors declare that they have no conflicts of interest with the contents of this article.

## Abbreviations

CA: carbonic anhydrase.
DIDS: 4,4′-diisothiocyanatostilbene-2,2′-disulfonate.
IRMS: isotope ratio mass spectrometry.
NTA: nanoparticle tracking analysis.
pNPP: p-nitrophenyl phosphate.
STAS: sulfate transporter and anti-sigma factor antagonist.
VPDB: Vienna Pee Dee Belemnite.

---

**Figure S1.**
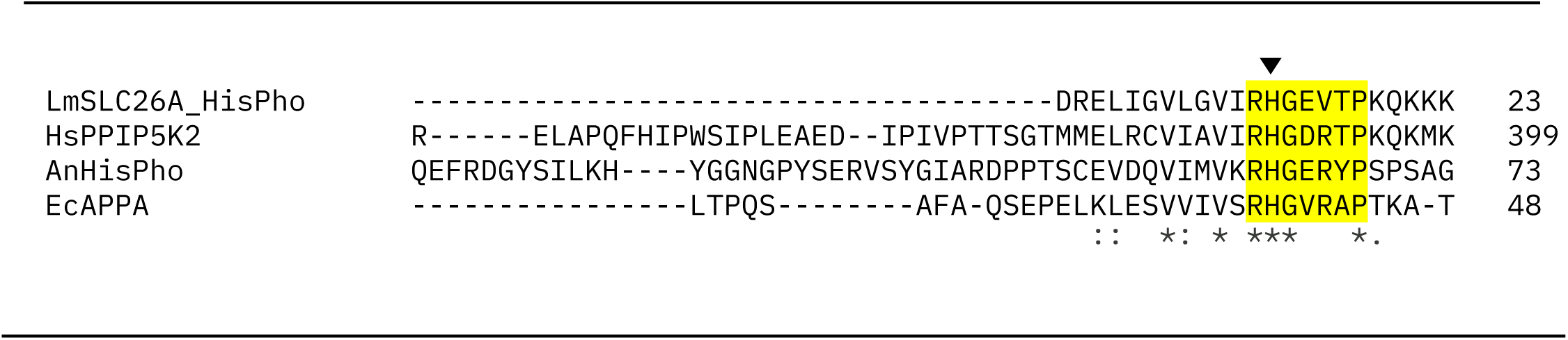
Conservation of the RHGXRXP motif in the predicted histidine phosphatase domain of LmSLC26A. Multiple sequence alignment of the predicted histidine phosphatase (HisPho) domain of LmSLC26A (residues 730-999), identified using InterPro, with corresponding histidine phosphatase domains from *Escherichia coli* (EcAPPA), *Homo sapiens* (HsPPIP5K2), and *Aspergillus niger* (AnHisPho). The conserved RHGXRXP catalytic signature motif characteristic of histidine phosphatases is highlighted in yellow. The catalytic histidine residue within the motif is indicated with black arrowhead. The RHGXRXP motif is largely conserved in LmSLC26A except the second arginine residue.

**Figure S2.**
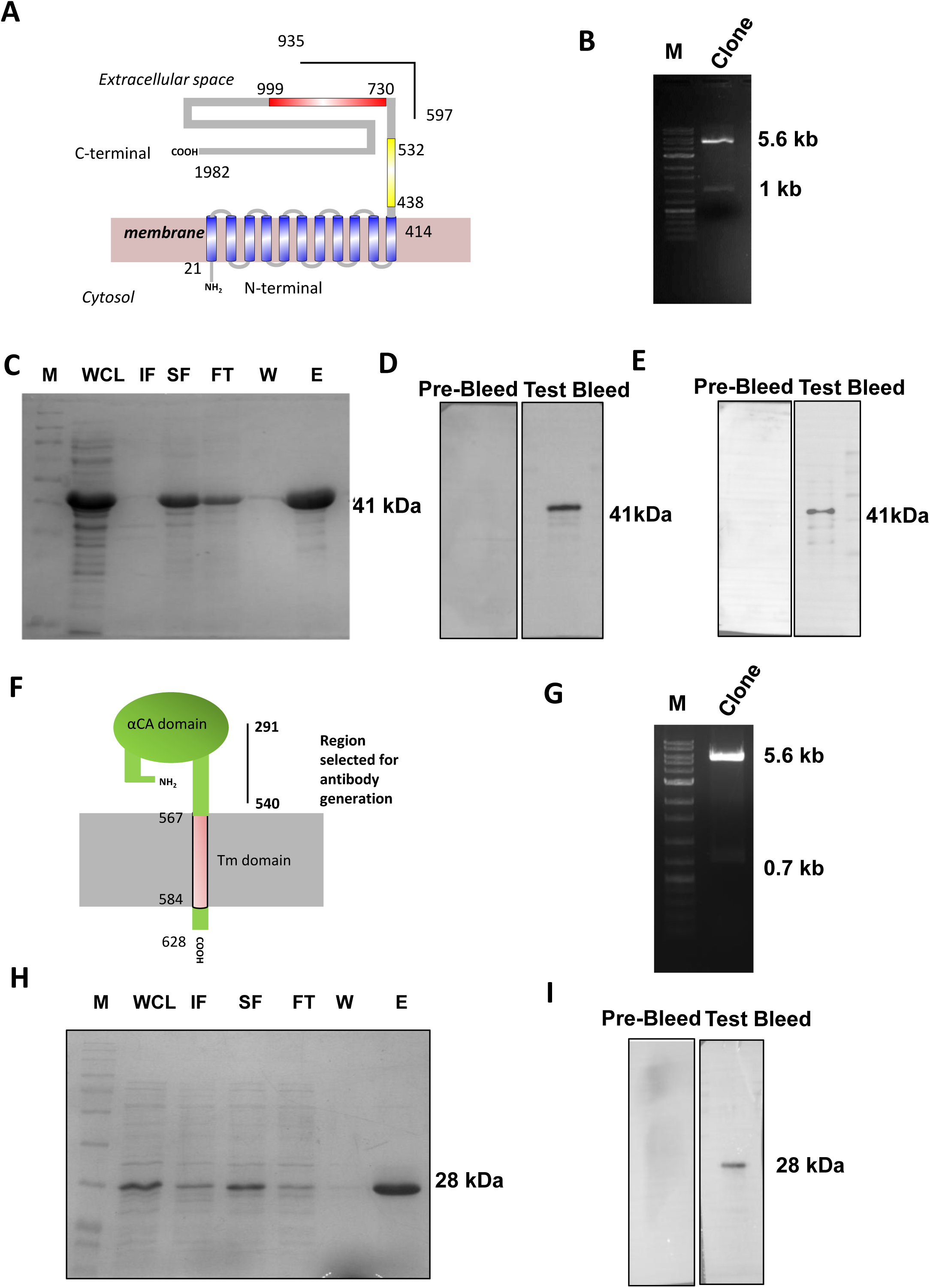
Generation and validation of antibodies against LmSLC26A and LmCA2. (A) Topological schematic of LmSLC26A showing the transmembrane domain (blue), STAS domain (yellow), and histidine phosphatase domain (red). The black line indicates the region selected for antibody generation. (B) Cloning of LmSLC26A^597-935^ into the pET-28a vector. Upon double digestion with EcoRI and HindIII, bands corresponding to the ∼1 kb insert and ∼5.6 kb vector backbone were observed. M, molecular weight marker. (C) Purification of LmSLC26A^597-935^ from a bacterial expression system using Ni-NTA affinity chromatography. WCL, whole-cell lysate; IF, insoluble fraction; SF, soluble fraction; FT, flow-through; W, wash; E, elution fraction. (D) Validation of LmSLC26A^597-935^ antibody generated in mice. Western blot of purified antigen probed with pre-bleed and immune sera. (E) Validation of LmSLC26A^597-935^ antibody generated in rabbit. Western blot of purified antigen probed with pre-bleed and immune sera. (F) Topological schematic of LmCA2 showing the transmembrane domain (pink). The black line indicates the region selected for antibody generation. (G) Cloning of LmCA2^291-540^ into the pET-28a vector. Double digestion with BamHI and EcoRI yielded bands corresponding to the ∼0.7 kb insert and ∼5.6 kb vector backbone. M, molecular weight marker. (H) Purification of LmCA2^291-540^ from a bacterial expression system using Ni-NTA affinity chromatography for antibody generation. WCL, whole-cell lysate; IF, insoluble fraction; SF, soluble fraction; FT, flow-through; W, wash; E, elution fraction. (I) Validation of LmCA2^291-540^ antibody generated in mice. Western blot of purified antigen probed with pre-bleed and immune sera.

**Figure S3.**
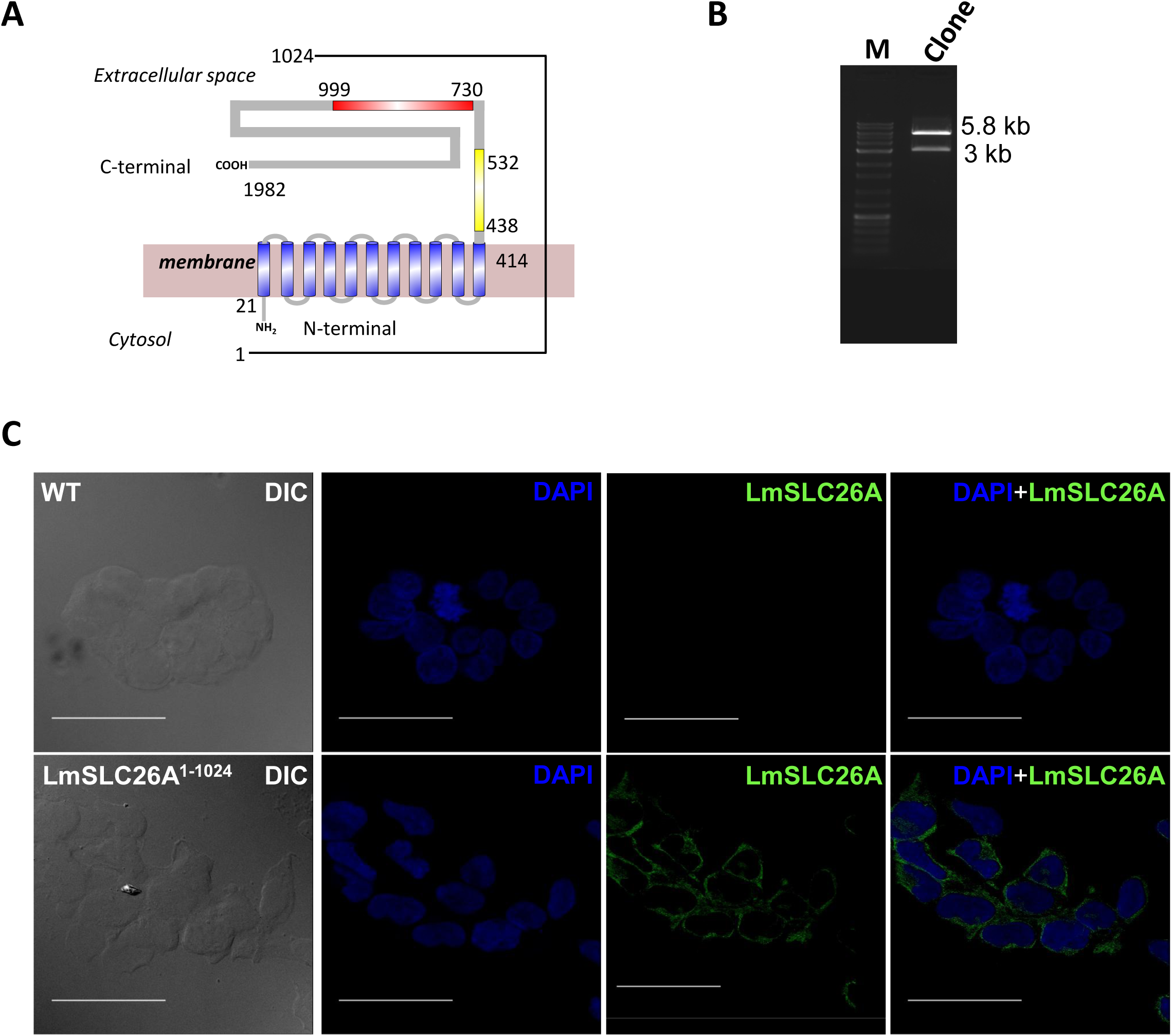
Validation of specificity of LmSLC26A antiserum by heterologous expression in HEK-293 cells. (A) Topological schematic of LmSLC26A showing the transmembrane domain (blue), STAS domain (yellow), and histidine phosphatase domain (red). The region corresponding to amino acids 1-1024 used for heterologous expression is indicated. (B) Cloning of LmSLC26A^1-1024^ into the pCXN vector. Upon digestion with EcoRI, bands corresponding to the ∼3 kb insert and ∼5.8 kb vector backbone were observed. M, molecular weight marker. (C) Stable HEK-293 cells expressing LmSLC26A1-1024 (bottom row) and Wild-type HEK-293 cells (top row) were immunostained with anti-LmSLC26A (green). DAPI (blue) marks the nucleus. DIC images (left) are shown alongside corresponding immunofluorescence images. Cells were imaged using a Leica SP8 confocal microscope. Scale bar: 50 μm.

**Figure S4.**
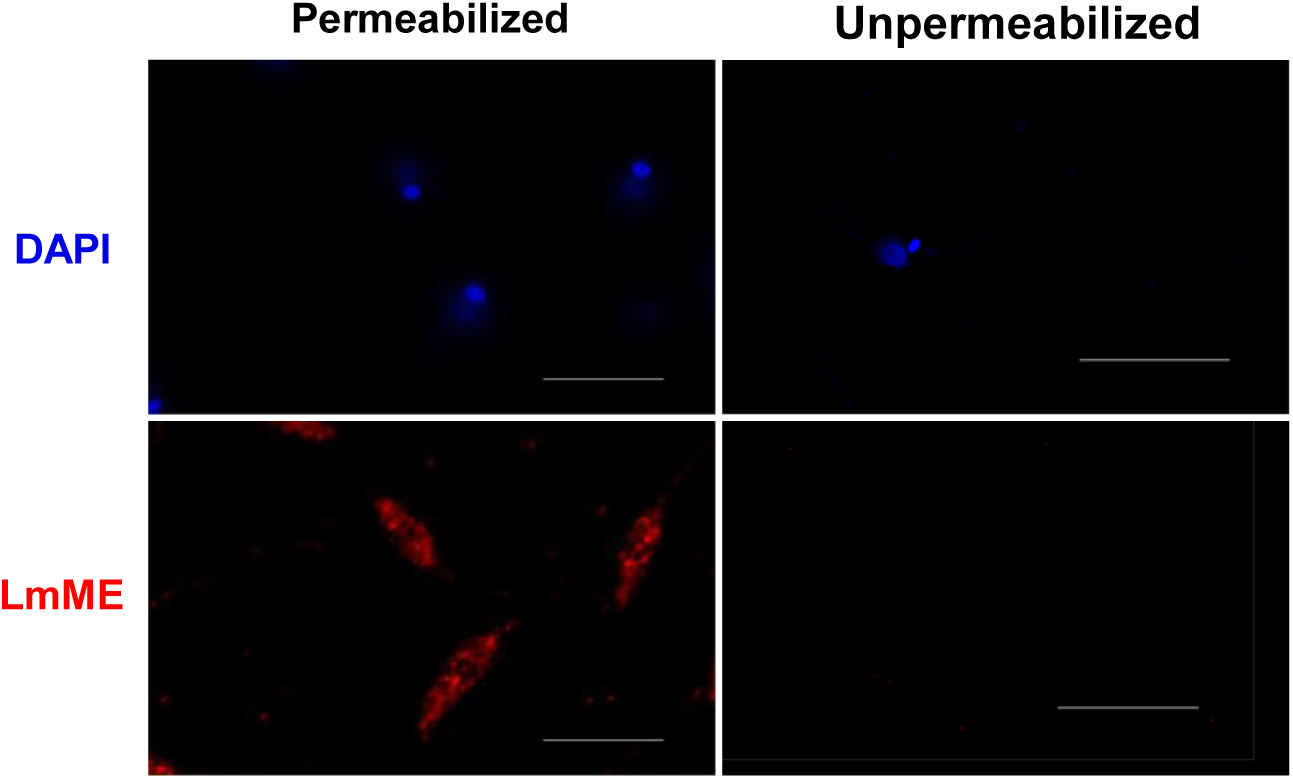
Immunofluorescence staining of *L. major* malic enzyme under permeabilized and non-permeabilized conditions. *L. major* cells were immunostained for the intracellular protein malic enzyme (LmME). Signal was detected under permeabilized conditions (left panels), whereas no signal was observed in non-permeabilized cells (right panels). DAPI was used to stain the nucleus and kinetoplast. Scale bar: 10 μm.

**Figure S5.**
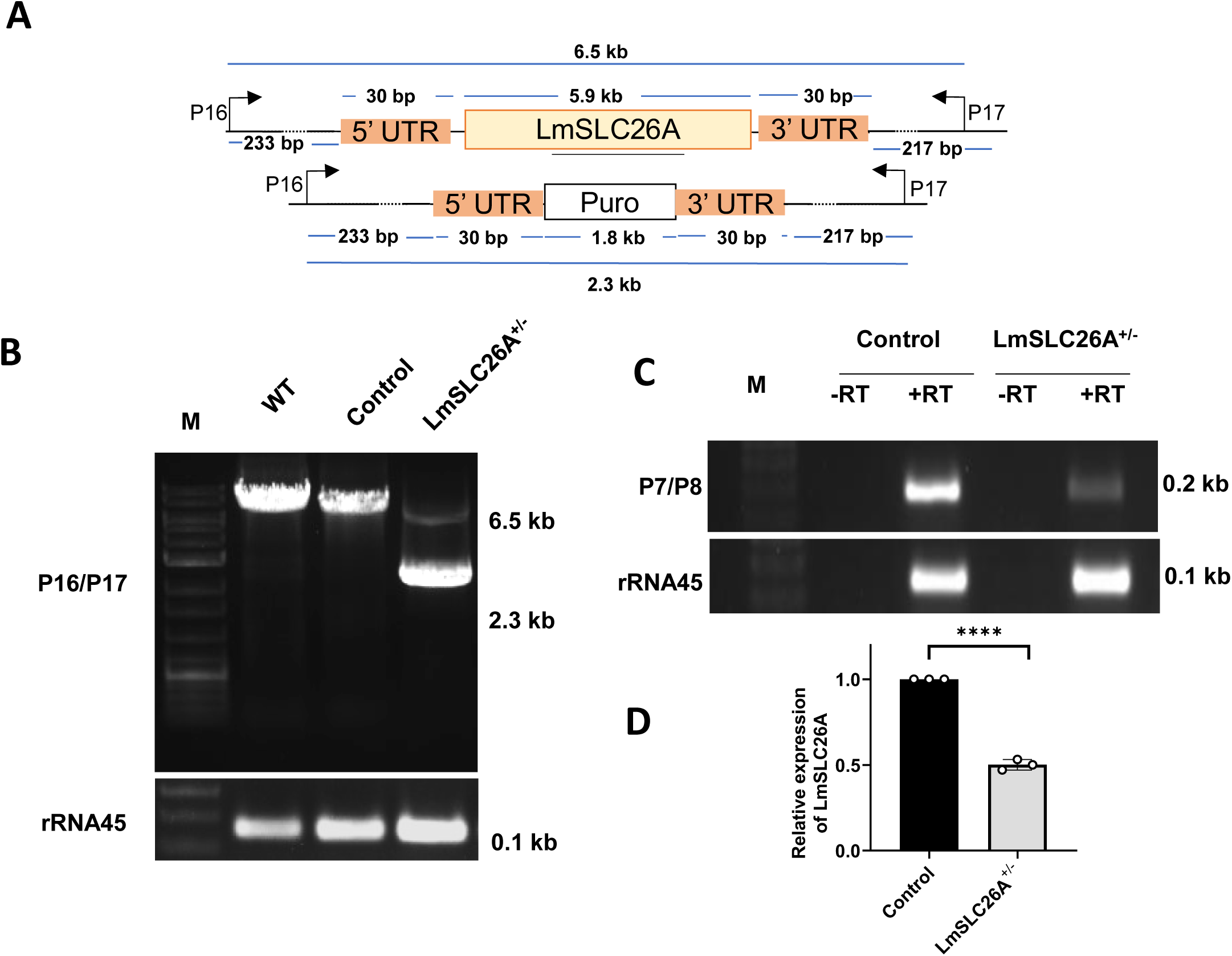
Verification of LmSLC26A^+/-^ at genomic and RNA level (A) Schematic representation of the genomic locus of LmSLC26A^+/-^ cells. (B) PCR analysis using gene flanking primers P16/P17. A ∼6.5 kb band was observed in WT, control (LmCas9:T7Pol), and LmSLC26A^+/-^ cells, while an additional ∼2.3 kb band was detected only in LmSLC26A^+/-^ cells. The lower panel shows amplification of rRNA45 as a loading control. (C) Comparison of LmSLC26A expression at the RNA level. Total RNA was isolated from *L. major* cells, treated with DNase, and reverse-transcribed to cDNA, followed by amplification using internal primers P7/P8. “+RT” and “−RT” indicate reactions performed in the presence and absence of reverse transcriptase, respectively. (D) Densitometric analysis of LmSLC26A transcript levels. Error bars represent SD from three independent experiments. ****P = 0.0000093 (Student’s t-test).

**Figure S6.**
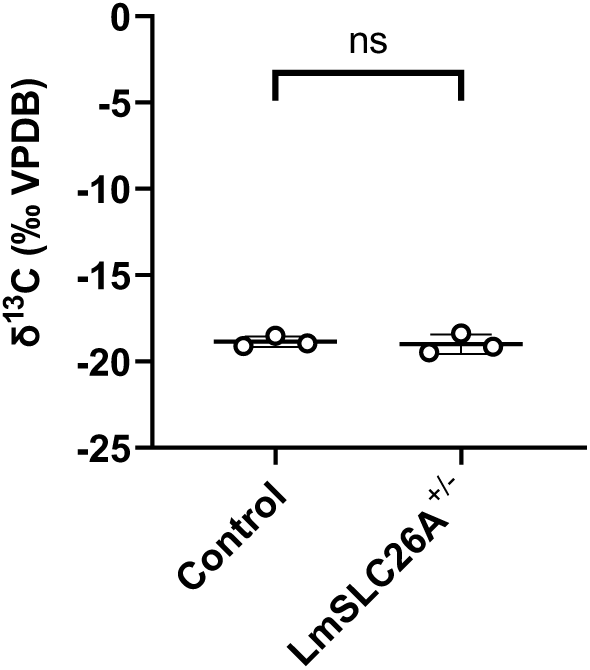
Bar graph showing basal δ^13^C values of control (LmCas9:T7Pol) and LmSLC26A^+/-^ strains, measured prior to addition of ^13^C-bicarbonate using IRMS. Data are expressed ‰ VPDB values. Error bars represent SD from three independent experiments. ns, not significant (Student’s t-test).

**Table S1.** Percentage Identity of LmSLC26A with different human bicarbonate transporters*.

| **Protein family** | **Protein aligned with** | **Percentage identity with LmSLC26A (in %)** | **Percentage similarity with LmSLC26A (in %)** |
|---|---|---|---|
| SLC26 | SLC26A1 | 25 | 44 |
|  | SLC26A2 | 24 | 42 |
|  | SLC26A3 | 24 | 46 |
|  | SLC26A4 | 25 | 47 |
|  | SLC26A5 | 27 | 45 |
|  | SLC26A6 | 27 | 47 |
|  | SLC26A7 | 25 | 45 |
|  | SLC26A8 | 22 | 42 |
|  | SLC26A9 | 24 | 43 |
|  | SLC26A10 | 25 | 45 |
|  | SLC26A11 | 32 | 50 |
| SLC4 | SLC4A1 | No significant match found | No significant match found |
|  | SLC4A2 |  |  |
|  | SLC4A3 |  |  |
|  | SLC4A4 |  |  |
|  | SLC4A5 |  |  |
|  | SLC4A7 |  |  |
|  | SLC4A8 |  |  |
|  | SLC4A9 |  |  |
|  | SLC4A10 |  |  |
|  | SLC4A11 |  |  |

<sup>\*</sup>Percentage identity and similarity determined using NCBI BLAST

**Table S2.**
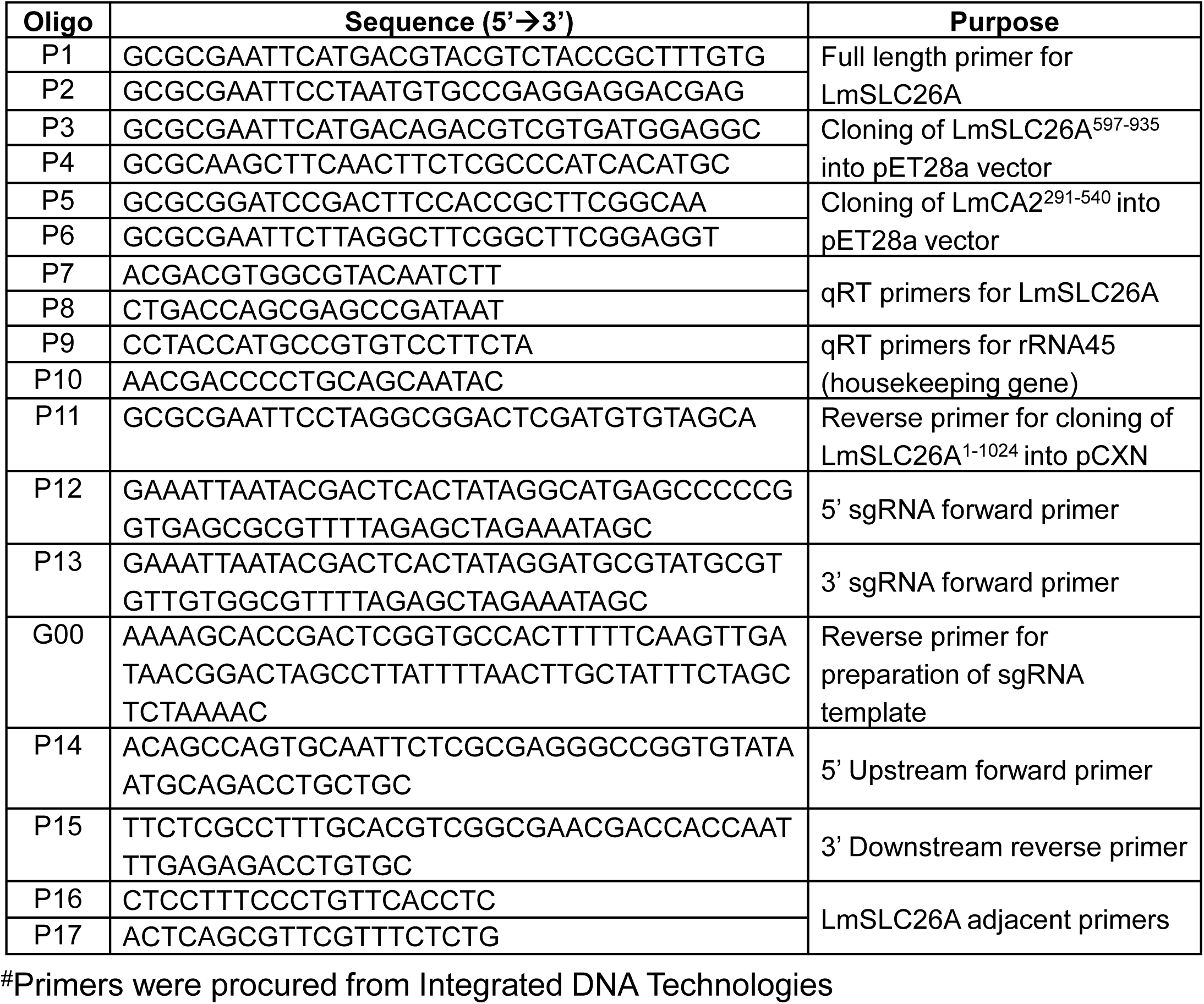
Oligonucleotides used in this paper^#^.

**Table S3.** Antibodies used in this paper.

| **Antibody** | **Host** | **IF (dilution)** | **WB (dilution)** | **Source** | **Catalogue number/ Reference** |
|---|---|---|---|---|---|
| Anti-LmSLC26A | Mouse | 1:200 | 1:1000 | In this paper | - |
| Anti-LmSLC26A | Rabbit | 1:50 | - | In this paper | - |
| Anti-LmCA2 | Rabbit | 1:200 | 1:1000 | In this paper | - |
| Anti-LmME | Rabbit | 1:200 | - | Mondal et al, 2021 | - |
| Anti-Rabbit IgG Peroxidase Goat | Goat | - | 1:4000 | Sigma Aldrich | A9169 |
| Anti-Mouse IgG Peroxidase Goat | Goat | - | 1:4000 | Sigma Aldrich | A9044 |
| Goat Anti-Rabbit Alexa-488 | Goat | 1:800 | - | Thermo Scientific | A11008 |
| Goat Anti-Rabbit Alexa-568 | Goat | 1:800 | - | Thermo Scientific | A11011 |
| Goat Anti-Mouse Alexa-488 | Goat | 1:800 | - | Thermo Scientific | A11001 |
| Goat Anti-Mouse Alexa-568 | Goat | 1:800 | - | Thermo Scientific | A11031 |

## Notes

### Competing Interest Statement

The authors have declared no competing interest.

### Summary of Updates

The tiltle has been slightly changed and new data has been added

